# Novel tunable spatio-temporal patterns from a simple genetic oscillator circuit

**DOI:** 10.1101/2020.07.06.190199

**Authors:** Guillermo Yáñez Feliú, Gonzalo Vidal, Macarena Muñoz Silva, Timothy J. Rudge

## Abstract

Multicellularity, the coordinated collective behaviour of cell populations, gives rise to the emergence of self-organized phenomena at many different spatio-temporal scales. At the genetic scale, oscillators are ubiquitous in regulation of multicellular systems, including during their development and regeneration. Synthetic biologists have successfully created simple synthetic genetic circuits that produce oscillations in single cells. Studying and engineering synthetic oscillators in a multicellular chassis can therefore give us valuable insights into how simple genetic circuits can encode complex multicellular behaviours at different scales. Here we develop a study of the coupling between the repressilator synthetic genetic ring oscillator and constraints on cell growth in colonies. We show *in silico* how mechanical constraints generate characteristic patterns of growth rate inhomogeneity in growing cell colonies. Next, we develop a simple one-dimensional model which predicts that coupling the repressilator to this pattern of growth rate via protein dilution generates travelling waves of gene expression. We show that the dynamics of these spatio-temporal patterns are determined by two parameters; the protein degradation and maximum expression rates of the repressors. We derive simple relations between these parameters and the key characteristics of the travelling wave patterns: firstly, wave speed is determined by protein degradation and secondly, wavelength is determined by maximum gene expression rate. Our analytical predictions and numerical results were in close quantitative agreement with detailed individual based simulations of growing cell colonies. Confirming published experimental results we also found that static ring patterns occur when protein stability is high. Our results show that this pattern can be induced simply by growth rate dilution and does not require transition to stationary phase as previously suggested. Our method generalizes easily to other genetic circuit architectures thus providing a framework for multi-scale rational design of spatio-temporal patterns from genetic circuits. We use this method to generate testable predictions for the synthetic biology design-build-test-learn cycle.

## 1 Introduction

Multicellularity and collective cell behaviour exemplify the emergence of complex patterns and structures across scales in living systems. When cells interact they can generate higher order patterns of gene expression (differentiation) as well as patterns of mechanical stresses and strains [1, 2]. This process takes place in natural phenomena such as embryonic development, tumor formation, wound healing, among others [3, 4, 5, 6, 7]. Understanding how these patterns are generated and maintained will enable applications in tissue engineering and regenerative medicine. However, natural emergent multicellular phenomena present numerous unknown processes that pose difficulties for understanding the fundamental mechanisms underlying pattern formation.

Synthetic biology applies design principles to generate combinations of genetic parts that perform a given function, for example oscillation, which at the same time helps us to understand the complexity inherent to natural systems. The prototypical engineering process is the design-build-test-learn cycle, which is an iterative process relying heavily on models of genetic circuit function. A variety of genetic circuits have been designed, analysed, simulated and then implemented in this way. These synthetic circuits simplify biological systems reproducing a specific function [8] such as toggle switches [9, 10], oscillators [11, 12, 13, 14], logic gates [15, 16, 17, 18] and arithmetic operators [19, 20].

While these circuits are often studied as dynamical systems in single cells or well mixed populations, the function of genetic circuits has also been studied in cell colonies [21, 3] through the engineering of patterns of gene expression such as symmetry breaking [22], Turing patterns [23], fractal patterns [24], tissue like strucures [25, 26], among others. These emergent spatio-temporal patterns depend on mechanical constraints on cells, which are the result of cell-cell and cell-substrate interactions. Thus, synthetic gene circuits can be engineered to generate higher order spatio-temporal patterns when coupled to mechanical constraints.

We focus here on the repressilator [11], a gene network that encodes a ring oscillator topology consisting of three repressors, where repressor 1 inhibits repressor 2, repressor 2 inhibits repressor 3 and repressor 3 inhibits repressor 1 (figure 1). In the original realization of this circuit topology [11] the circuit was subject to significant effects of noise and oscillations quickly became desynchronized. Recently, the circuit was revisited by [14] with microfluidics systems that allowed them to observe single cells oscillating synchronously in chemostatic conditions for long periods of time. In this work sources of noise were reduced in several ways. Firstly, the fluorescent reporters were integrated into the same low-copy plasmid as the repressilator reducing the standard deviation in amplitude greatly. They also removed the degradation tags and used a protease deletion strain (ΔclpXP) as the chassis to remove noise from enzymatic queuing [27, 28]. They also increased the effective repression threshold with a high-copy titration ‘sponge’ plasmid that sequesters a proportion of the TetR repressor, since low repression thresholds imply sensitivity to noisy repressor expression levels. These modifications allowed regular and sustained synchronous oscillations that peaked around every 14 generations. The circuit oscillated for approximately 18 periods before accumulating half a period of drift, demonstrating that cells remained synchronized for more than 250 generations. Strikingly, [14] were able to observe whole flasks of liquid bacterial culture oscillate synchronously, and bacterial colonies form coherent ring patterns at macroscopic scale. These findings show that the repressilator can be effectively isolated from noise, function in a robust and synchronous fashion, and is capable of forming spatial patterns.

**Figure 1:**
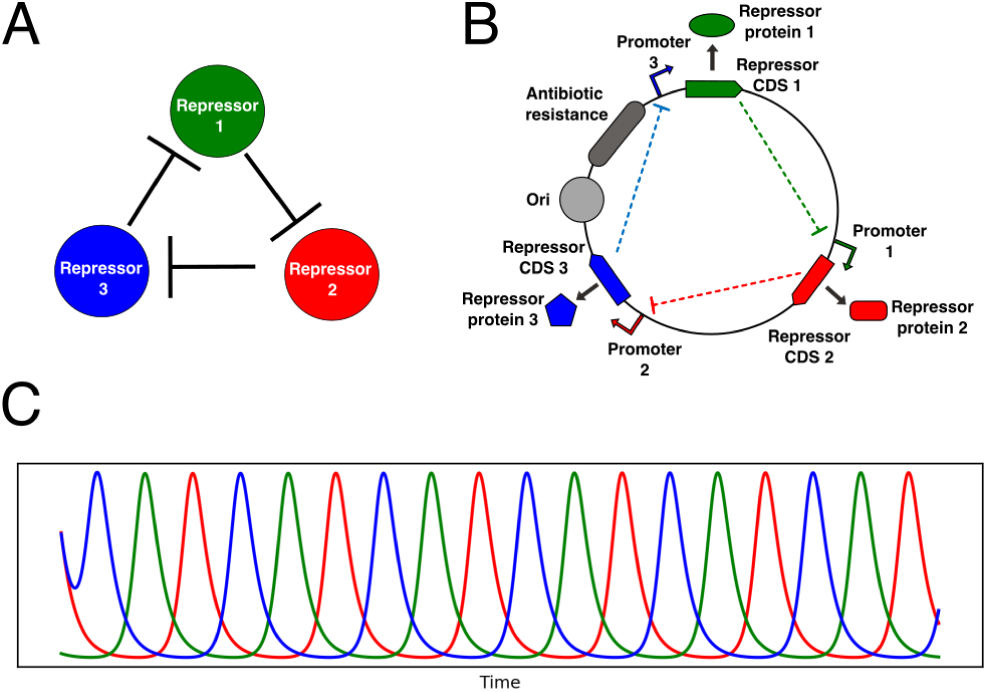
Repressilator genetic oscillator circuit. **(A)** Network representation of the repressilator, in the schematic 1 represses 2, 2 represses 3 and 3 represses 1. **(B)** Genetic circuit diagram of a plasmid encoding the repressilator. **(C)** Oscillating repressor concentrations over time computed by solving equation 11 with fixed growth rate. CDS 1, 2, and 3 are the complementary DNA sequences coding for each repressor, and Ori is the origin of replication.

Models are essential in the design process, they allow engineers to screen the parameter space looking for possible functional constructions [29, 30]. Synthetic biology has gone from intracellular dynamic models using ODEs [9, 11], and SSA [14, 23] to sophisticated collective behaviour models based on individual agents [31, 32] and integrated circuit-host models [33]. Using cellular scale individual-based models (IBMs) gives rich information about the emergent collective properties of cell populations due to the interactions between themselves and their environment. These models track cell growth and gene expression in ways analogous to experiments performed in controlled environmental conditions with specified properties such as viscosity, chemical concentrations, etc. This makes them an essential tool in the analysis and design of emergent properties of genetic circuits operating in multicellular chassis. However these models are complex and require significant computation time, highlighting the need for simple tractable mathematical and computational methods.

In this study we uncover novel spatio-temporal patterns of gene expression generated by the repressilator in growing cell colonies, and establish a simple method for their design. Since it is generalizable, this work provides a quantitative framework for multi-scale rational design of spatio-temporal patterns from genetic circuits. We provide testable predictions for the synthetic biology design-build-test-learn cycle for engineering repressilator spatio-temporal pattern.

## 2 Results

### 2.1 Growth rate variation in growing microcolonies

We consider the case of *Escherichia coli*, the cellular chassis for which the repressilator (figure 1) was designed, growing on a viscous substrate such as a hydrogel or PDMS (polydimethylsiloxane) and supplied with fresh nutrients via microfluidic channels. Growth is constrained by forces between cells and between cells and the substrate due to viscous drag [32]. The cells in such a system are constrained to a monolayer and form a quasi-two-dimensional array of extending rod shapes [34, 35]. We used an individual-based model [32] to characterize the distribution of cell growth rates across a two dimensional cell monolayer growing in such conditions over time. We simulated the growth of microcolonies from single cells to populations of approximately 60,000 cells with a radius of approximately 200 cell diameters. Figure 2A-C shows the development of a colony from approximately 5000 cells to 50,000 cells. The distribution of growth rates across the colony has a clear radially symmetric pattern with a maximum at the edge of the colony (Supplementary Material, figure S1). This is as expected [36, 37] since the cells at the edge of the colony are relatively unconstrained. Thus individual bacteria inside microcolonies perceive a different biophysical environment depending on their spatial position. Taking radial averages of growth rate on growing colonies over a range of time points we see the same exponential decay relative to the colony edge (figure 2E), leading to a simple model for the cell growth rate as a function of the radial position of the cell with respect to the colony edge *r*(*t*),

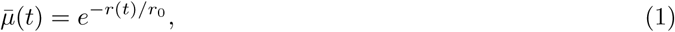

where *r*_0_ (8.23 ± 1.69 cell diameters) is the characteristic length scale of the radial variation in growth rate, and we have normalized by *µ*_0_ – the maximal unconstrained growth rate of the cells. These results suggest that growth rate time dynamics are determined by radial distance from the colony edge. After a short transient, the colony edge moves with constant velocity *v*_*front*_ = 5.00 so that *R*_*max*_ increases linearly (figure 2F).

**Figure 2:**
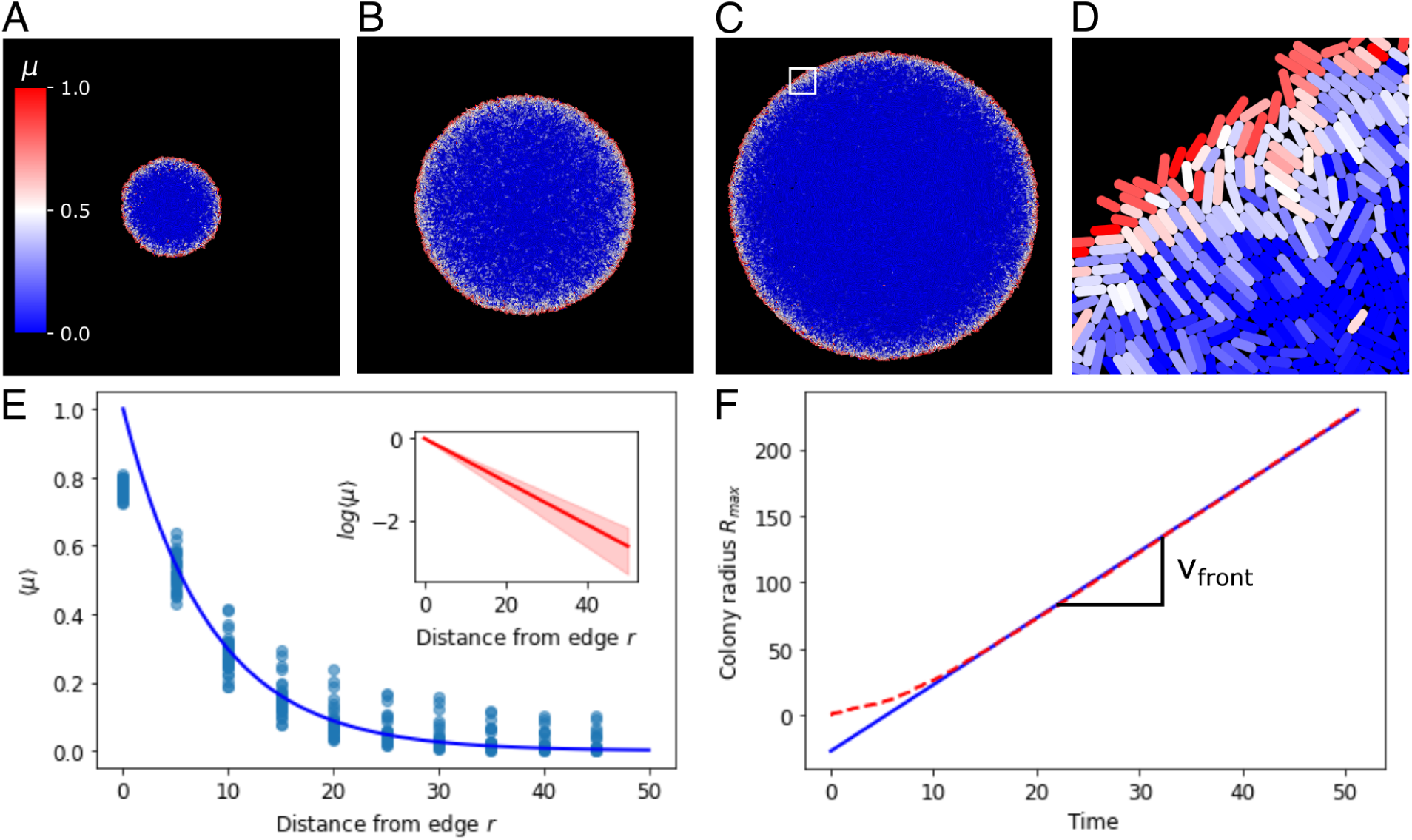
Average growth rate decays exponentially with distance relative to the edge. **(A), (B)** and **(C)** are growing colonies at 5000, 25,000 and 50,000 cells respectively. **(D)** is a zoom from the white square region in C. Single cells are colored according to their growth rate, which ranges between 0 and 1. **(E)** Average growth rate ⟨*µ*⟩ at different distances from the colony edge, blue line is an exponential function (*e*^−*r/r*0^) with *r*0 = 8.23 fitted to the binned data (blue dots). Inset shows the log scale of the average growth rate ⟨*µ*⟩ at different distances from the edge. **(F)** Colony radius (*R*_*max*_) over time (dashed red line), linear fit to data at t>10 giving *v*_*front*_ = 5.00 the velocity of the colony edge (blue line).

Assuming a two-dimensional densely packed monolayer with random cell orientations, growth is isotropic and expansion is equal in all directions. The area expansion rate approximates the growth rate and is given by the divergence of the velocity field,

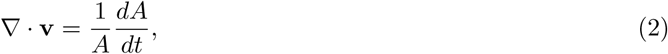

where *A* is the cell area and **v** is the velocity. Since growth is isotropic we may decompose the expansion rate equally into its radial and perpendicular components. Considering the velocity *v* in the radial direction *r*, and velocity *w* in the perpendicular direction *s*, and expanding the divergence term, equation 2 gives,

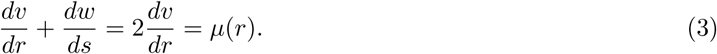

Hence, with *v* = *dr/dt*, and considering only the radial direction, we can rescale time as *t* → *tµ*_0_ and radial distance as *r* → *r/r*_0_ to obtain,

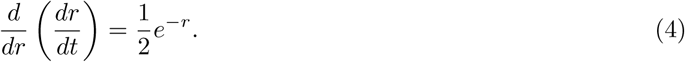

Integrating by *r* and *t* results in,

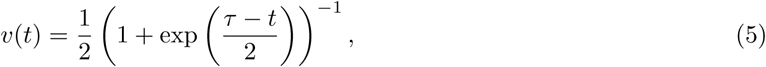

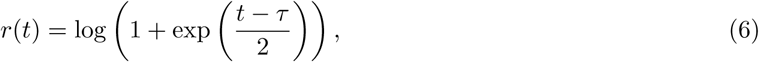

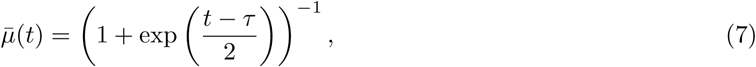

where *τ* = −2log(exp(*r*(0)) − 1), and *r*(0) is the initial radial position of the cell.

Equations 5-7 give us valuable insights into the system behaviour (figure 3). The velocity in the radial direction (away from the colony edge) is a sigmoidal logistic function and saturates to a velocity of *v* = 1*/*2 as *r* increases (figure 3A). This gives the front velocity as *v*_*front*_ = 1*/*2 in rescaled units. Correspondingly the radial position relative to the colony edge *r* tends towards linear increase at velocity *v* = 1*/*2 as the cell becomes effectively stationary relative to the colony center (figure 3B). In real units this means that the front velocity is *v*_*front*_ = *r*_0_*/*2, where *r*_0_ is the length scale of growth rate variation (equation 1). This is consistent with our individual based simulations (figure 2F), in which *v*_*front*_ = 5.00 and *r*_0_ = 8.23 ± 1.69. The growth rate is also sigmoidal and tends from maximum at the colony edge to zero as the cell moves away from the growing front of the colony (figure 3C).

**Figure 3:**
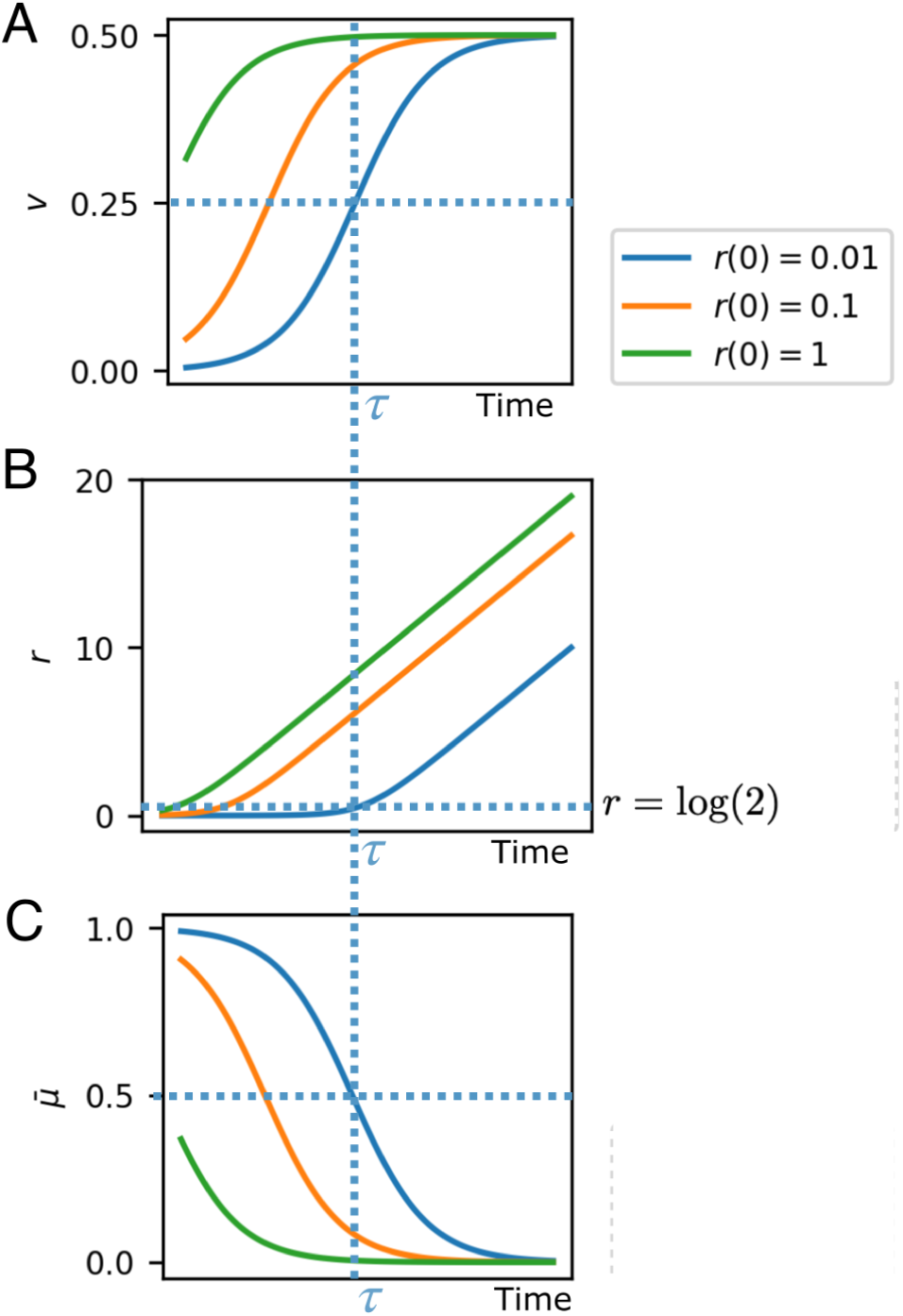
Analytical solutions for cell velocity (A) and position (B) relative to the colony edge, and growth rate (C), based on exponential decay of growth rate with distance from colony edge. At critical time *τ* the system switches between two regimes: high growth/low velocity, and low growth/high velocity.

The critical time *τ*, depending on the initial cell position, is the time at which the growth rate and velocity are at their half maximum values, and the cell is at radial position *r* = log(2) (figure 3, dashed lines). At this time the cell switches from a high growth, low velocity regime (remaining close to the colony edge), to a high velocity, low growth regime (remaining stationary while the colony edge propagates). The time *τ* for this switch to occur is greater for cells closer to the colony edge, that is they remain in the fast growing regime for longer. Thus cells effectively experience a switch in growth rate and velocity at their critical time *τ*, which depends on their initial radial position in the colony.

### 2.2 Dynamic growth rate dependent mathematical model of the repressilator

Here we develop a simple mathematical model coupling the repressilator genetic circuit to growth rate variation via simple dilution of proteins. The repressilator can be considered as an abstract genetic circuit topology. We consider an implementation of this topology following the design modifications made by [14], which essentially isolated the circuit from noise and allowed sustained and synchronous oscillations over time scales up to 250 generations. Stochastic simulations performed with relevant parameters reproduced this behaviour, showing essentially deterministic behaviour (Supplementary figure S2), therefore we may use a simpler differential equation model to track the repressor concentration of each cell over time. A simple two-step model of the balanced repressilator genetic circuit (figure 1), a type of genetic ring oscillator, can be formulated as follows,

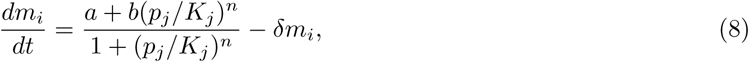

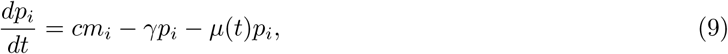

where *i* is one of the three genes, *j* is its corresponding repressor gene, *m*_*i*_, *p*_*i*_ are the mRNA and protein concentrations respectively, *a* is the constitutive transcription rate, *b* is the leaky or repressed transcription rate, *µ* is the instantaneous growth rate of the cell or population of cells, *γ* is the protein degradation rate, and *δ* is the mRNA degradation rate. Order of magnitude estimates of these parameters are given in Supplementary Material.

Since mRNAs are typically short lived (see Supplementary Material), we may assume quasi-steady state concentrations and the system is,

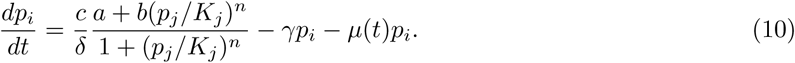

Rescaling protein concentration as *p*_*j*_ →*p*_*j*_*/K*_*j*_ and time by *t* → *tµ*_0_ with *µ*_0_ the maximal growth rate, assuming that basal expression is negligible, and combining with equation 7,

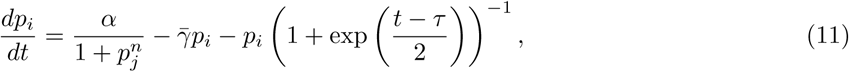

where *α* = *ca/δµ*_0_*K* (order of magnitude 10^4^, see Supplementary Material) is the steady state maximal gene expression rate and the constant 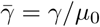 is the protein degradation rate as a fraction of the maximal growth rate (order of magnitude 1). This model depends on three dimensionless parameters *α*, 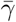 and *n*.

In this model we assume that the dominant effect of growth rate variation is by dilution of proteins. While there is evidence for growth rate dependencies of transcription and translation rates and plasmid copy number [38, 39, 40], all of which affect the parameters of the model, these effects have only been observed due to different biochemical environments. In spatially constrained cell populations the shape of the growth profile 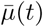 depends both upon the biochemical environment and mechanical constraints [41, 42, 43, 24, 34, 37, 44]. At the typical bacterial microcolony scale the biochemical environment is essentially uniform in space due to the fast diffusion of small molecules like sugars and aminoacids [45, 46, 47]. Using microfluidic devices cells can be maintained in constant fresh media allowing observation of the long term dynamics of genetic circuits [13, 48, 14]. Under these conditions then the predominant factors determining growth rate are physical forces and constraints.

### 2.3 Protein dilution enables the repressilator to produce traveling waves in growing microcolonies

The model presented above (equations 5-7 and 11) describes the trajectories of cells as they move in the radial direction and change their protein concentrations over time. Assuming that the motion and growth of cells is not affected by the expression of repressor genes, this model describes the mean behaviour of cells starting from some initial radial position. It is obvious from these equations that in the absence of growth the system can only produce plane wave, homogeneously synchronized oscillations. However, in the presence of growth we have an explicit relation between cell position and protein expression rate, enabling spatio-temporal pattern formation.

Since growth dilutes proteins the effective degradation rate of the repressors is 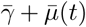. The effect of the sigmoidal growth rate switch on the repressilator is therefore to decrease the effective degradation rate of the repressors from 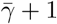 to 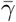 as cells move out of the growing regime (equation 7). [14] showed that increasing the degradation rate of repressors by appending a degradation tag reduced the period of oscillations *T*. This decrease was by approximately a factor of two at 37 C, with less effect at lower temperatures likely due to decrease in protease activity [49]. We confirmed this result using our model by numerically integrating equation 11 at fixed effective degradation rates 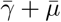 (figure 4). The frequency of oscillations *f* = 1*/T* was proportional to the degradation rate, with a slope depending on *α*, the maximum gene expression rate. Hence in colonies, as cells move away from the edge due to mechanical constraints the effective repressor degradation rate decreases and the period of their oscillations increases.

**Figure 4:**
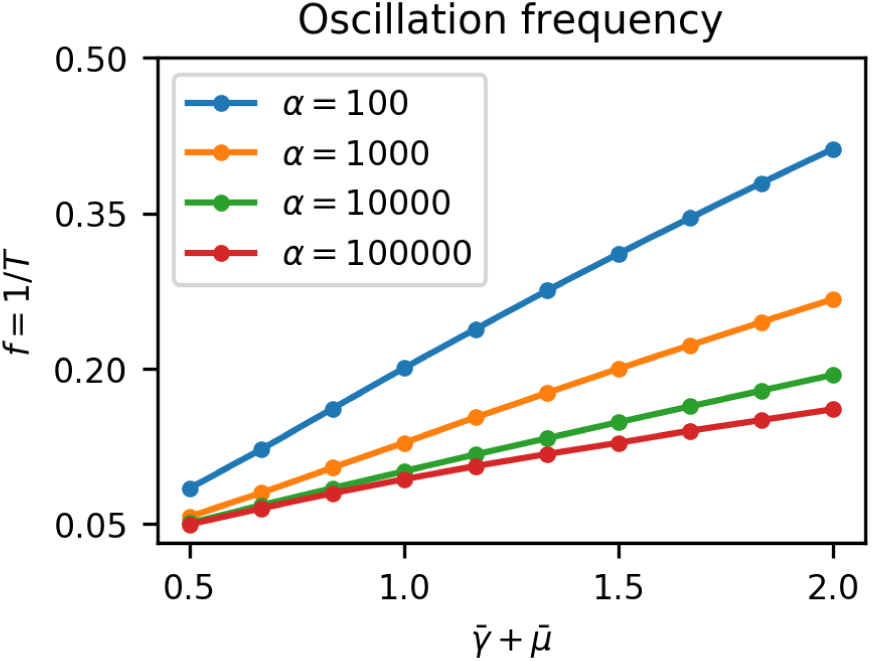
Oscillation frequency *f* (or period *T*) depends on the effective degradation rate of repressors 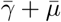. For a given maximum gene expression rate *α* the frequency is proportional to the effective degradation rate, and increasing *α* decreases the frequency.

After the critical time *τ* the period of oscillations increases as the cell switches from high growth rate and low velocity to low growth rate and high velocity (equations 5-7). This means that there is effectively an interior region oscillating with long period *T*_*int*_ and an exterior region oscillating with short period *T*_*ext*_. The phase offset between peaks of the two signals after some time Δ*t* is,

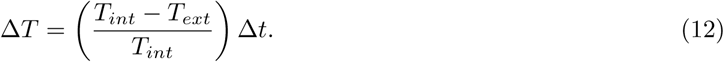

When the phase offset Δ*T* = *T*_*ext*_ we have in phase oscillations. The time required for a cell to achieve this phase offset is the time spent in the high growth regime *τ*, plus the time spent in the low growth regime, hence,

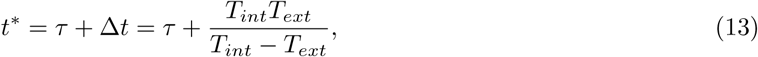

is the time at which the cell is in phase with the edge of the colony, the wave source. At time *t*\* the distance from the edge *r*(*t*\*) of this cell can be obtained from equation 6, and since this is the peak-to-peak distance it gives the wavelength *λ*. Assuming that exp(*t*\*) ≫ 1,

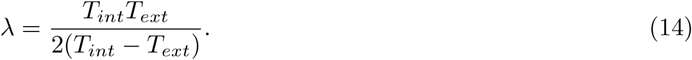

The wave propagation velocity is 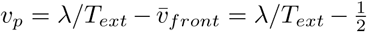, where 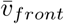 is the velocity of the colony edge in normalized distance units, hence,

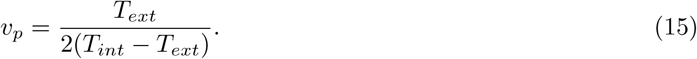

Equations 14 and 15 show that when *T*_*ext*_ *< T*_*int*_ the system generates traveling waves with finite wavelength and wave speed. When *T*_*int*_ = *T*_*ext*_, there is no effect of mechanical constraint on oscillation period and we find that *v*_*p*_ = ∞ and *λ* = ∞, meaning that the system forms homogeneous plane waves with the whole colony oscillating synchronously. As *T*_*int*_ → ∞ the interior does not oscillate and we find that *v*_*p*_ → 0 and *λ* → *T*_*ext*_*/*2 and we thus have static rings of gene expression with spatial wavelength 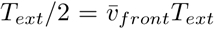. This is the case when protein degradation is negligible 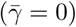, a condition under which the repressilator has been shown to form static rings in growing colonies [14]. Thus we have shown analytically that growth rate heterogeneity induces the repressilator to form either static rings or traveling waves in growing cell colonies, depending on the degradation rate of the repressors.

### 2.4 Novel spatio-temporal patterns emerging from repressilator dynamics

To test the predicted spatio-temporal patterns we integrated equation 11 from a range of initial cell radial positions to construct the kymograph *p*_*i*_(*R, t*), where *R* = *t/*2 − *r* is the rescaled distance from the center of the colony. A kymograph (figure 5A-C) represents the spatial dynamics of a one-dimensional system, such as ours, evolving over time. Each point in the kymograph represents the state of the system at a given time (x-axis) and position (y-axis) with a color. By taking radial averages the kymograph fully characterizes radially symmetric spatio-temporal patterns with the vertical axis representing distance from the center of the colony. Here we reflect the kymograph to represent the symmetric structure of the pattern (figure 5A-C), whereby the growth of the colony can be seen as two linearly expanding borders forming a triangular shape. The slope of this border is the front speed 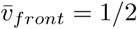. The color represents the radially averaged repressor protein concentrations (red, green, and blue) at each position at each time point, normalized to their maximum values. The corresponding predicted colony pattern is shown inset in figure 5C. Stripes in the kymograph represent rings of gene expression. Horizontal stripes show static rings since their radial position does not change (figure 5A). Vertical stripes would represent in-phase homogeneous oscillations since they do not vary in space (figure 5B). Diagonal stripes represent traveling waves, moving rings of repressor expression, since they vary in both space and time (figure 5C). Hence we confirm our theoretical prediction of traveling wave patterns.

**Figure 5:**
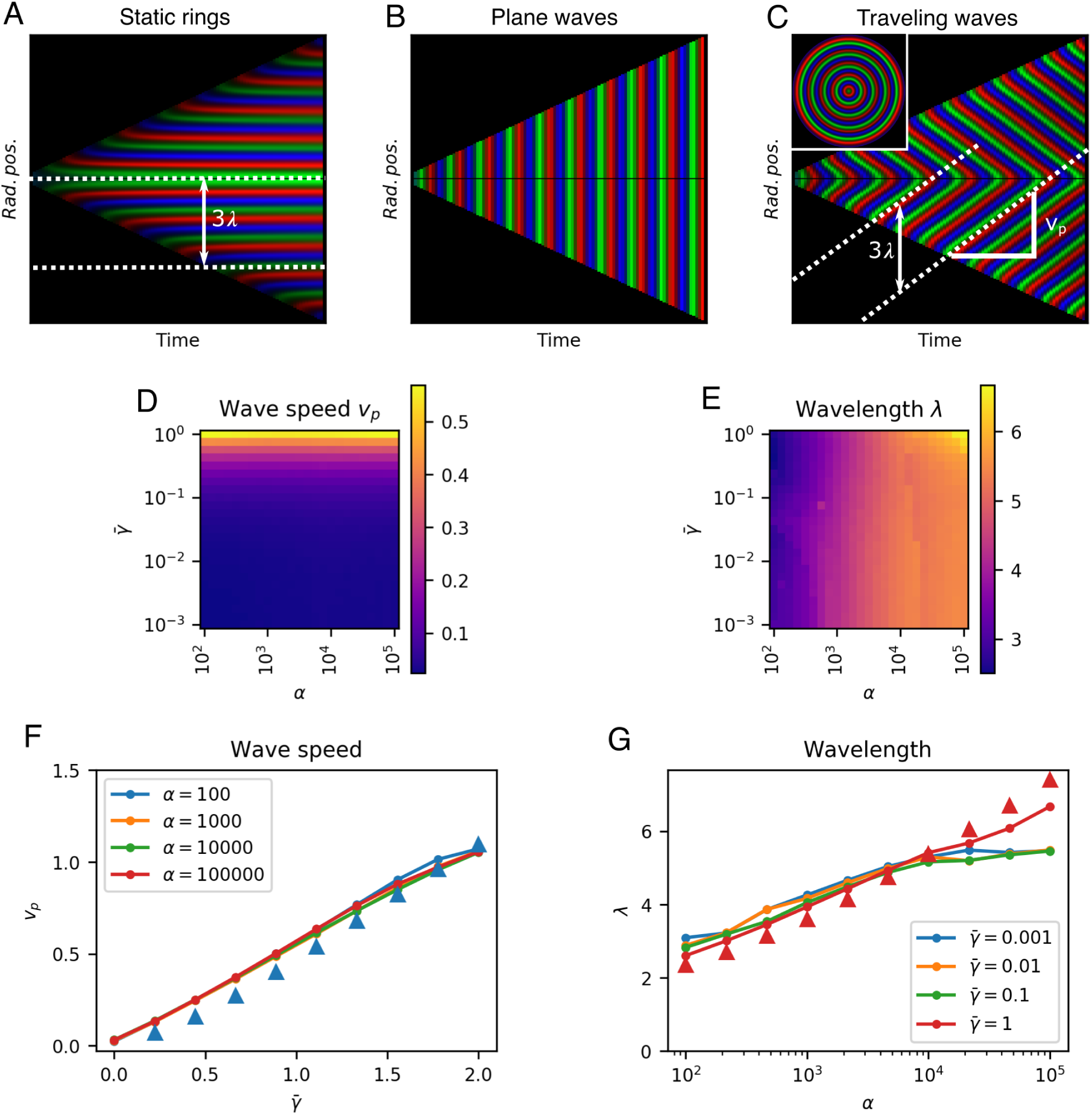
Parameter dependence characterization for 1D model. **(A-C)** Kymographs of growing colonies show protein concentrations as a function of radial position over time, with no protein degradation (**A**,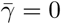, *α* = 10^4^) we obtain static rings, with no growth rate dilution (**B**, 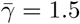, *α* = 10^4^) we see plane waves, and with growth rate dilution and protein degradation (**C**, 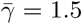, *α* = 10^4^) we have traveling waves. White dashed lines show wave trajectories. The distance between two trajectories is the wavelength *λ*, and the slope is the wave speed *v*_*p*_. Inset in **(C)** is the whole colony at the end of the experiment. **(D), (E)**. Heatmaps of wave speed *v* and wavelength *λ* respectively over a range of *α* and 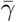. **(F)**. Effect of 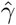 on wave speed *v* at different *α*. **(G)**. Effect of *α* on wavelength *λ* at different 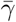. Triangles in **F** and **G** show analytical estimates for *α* = 100 and 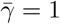 respectively.

The spatio-temporal dynamics of the system are described by two parameters that can be extracted from the kymographs. The slope of each stripe gives the wave speed *v*_*p*_, and the vertical peak-to-peak distance gives the spatial wavelength 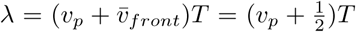, where *T* is the period of oscillations at the colony edge (the wave front) and 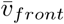 is the front velocity in normalized distance units (figure 5C).

### 2.5 Tuning the repressilator to control spatio-temporal pattern formation

In order to quantitatively test our theoretical predictions and to characterize the design space of these travelling wave patterns we scanned the parameter space within physiologically relevant ranges. We measured the wave speed *v*_*p*_ and wavelength *λ* of the system for 625 combinations of *α* and 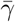 spanning four orders of magnitude to construct the phase space (figure 5D,E). We see that wavelength was predominantly determined by *α*, while wave speed depended on 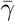. Travelling waves were observed for all values of *α* but clearly require non-zero protein degradation rate 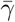 (figure 5F).

We observed that wave speed was proportional to 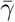, with 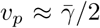 (linear fit 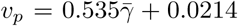). Static rings (*v*_*p*_ = 0) form when *γ* = 0. As 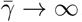, as is the case at growth arrest (*µ*_0_ = 0), we saw earlier that the system can only form plane waves with all parts of the colony oscillating in phase, which corresponds to *v*_*p*_ = ∞. Wavelength was predominantly but weakly affected by maximum gene expression rate *α* (figure 5G) following approximately *α* ≈ 10^*λ*−1^ (from the linear fit *λ* = 1.01 log10(*α*) + 0.998. These results are consistent with our theoretical predictions based on the oscillation period from equations 14-15 (figure 5F,G triangles).

We demonstrate the tuning parameters *α* and 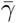 above using kymographs in figure 6. With no protein degradation 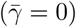 the system produces static rings of gene expression following the phase of each repressor (figure 6 A, B). In the kymograph this spatio-temporal pattern is observed as horizontal stripes of consecutive red, green, and blue, representing the three repressors. This confirms the observation of fixed ring patterns in colonies hosting a repressilator with stable repressor proteins [14]. The static ring patterns observed are therefore a special case of the more general travelling wave solution with velocity *v*_*p*_ = 0. These travelling waves are induced and modulated by protein degradation. In the intermediate case when protein degradation and growth are similar (figure 6 C, D, F) we see the clear emergence of a travelling wave solution. This spatio-temporal pattern is characterized by diagonal stripes in the kymograph, with steeper sloped lines indicating higher velocity of the waves (figure 5A). At lower protein degradation rate 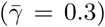 we see travelling waves with lower velocity (figure 6C, D). Hence protein degradation rate tunes the speed of the travelling waves. Changing *α* however does not affect the speed of the waves (figure 6A, C, E and B, D, F) but does change the wavelength of the travelling waves resulting in more spatial rings at lower *α* values. We note that increasing *α* also stabilizes the oscillations as found by [50, 14] (figure 6 panels below each kymograph).

**Figure 6:**
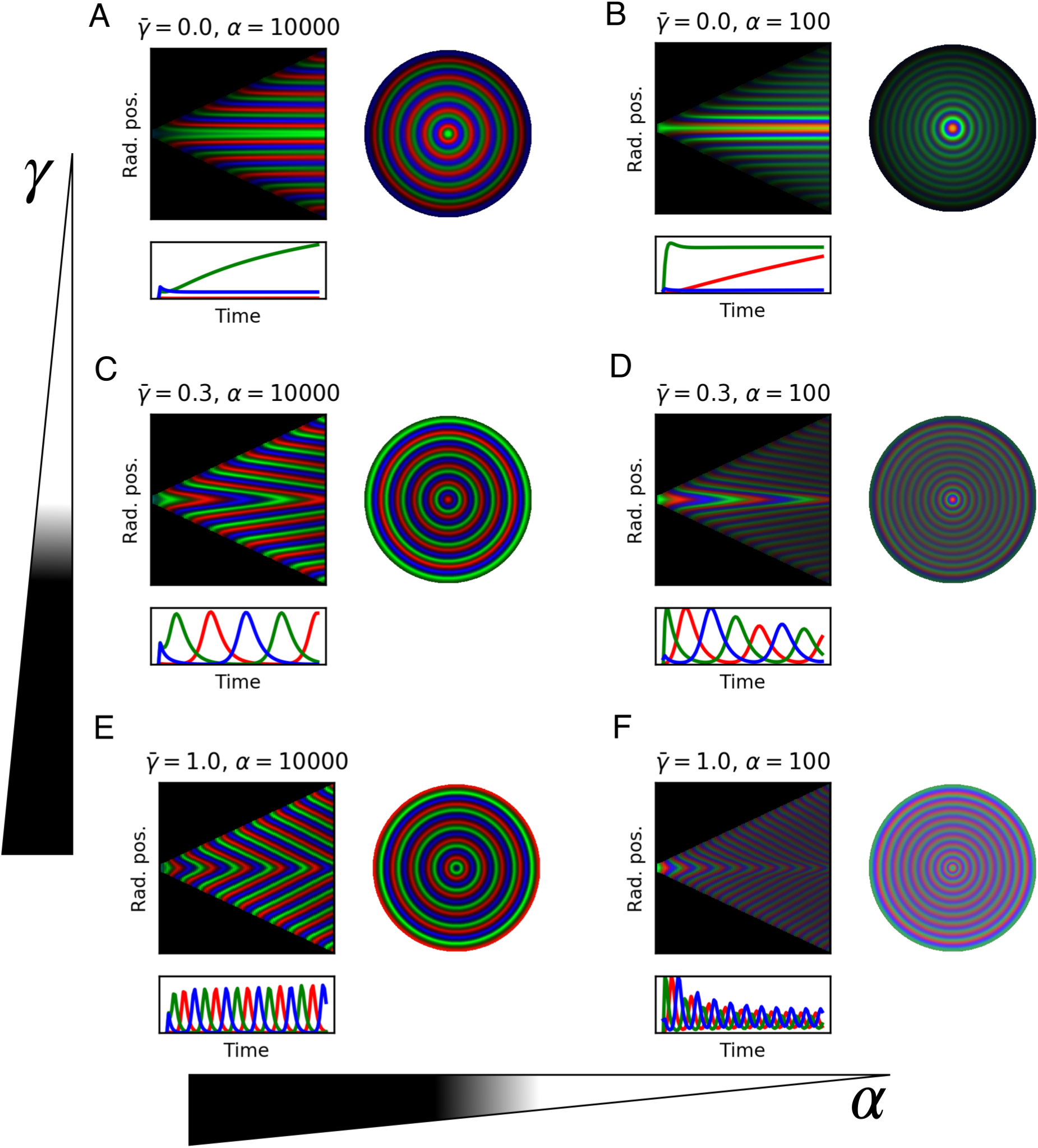
1D model simulations for selected parameters. Each panel shows: kymograph of the emerged pattern, where stripes show wave trajectories; trace of the three repressor concentrations over time in the center of the colony; the whole colony showing the final ring pattern. **(A)**. 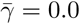, *α*=10.000. **(B)**. 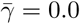, *α*=100. **(C)**. 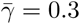, *α*=10.000. **(D)**. 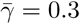, *α*=100. **(E)**. 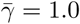, *α*=10.000. **(F)**. 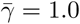, *α*=100.

### 2.6 Growing cell colonies generate travelling waves in quantitative agreement with predictions

To test if these predictions hold in constrained growing microcolonies of cells we used our individual based biophysical model of bacterial cell growth and division. We grew colonies from a single cell up to 60,000 cells, tracking each cell’s motion and protein expression levels according to equation 11 without the growth rate term (dilution was computed by the biophysical model). The results show, as predicted, the formation of symmetrical rings relative to the center of the colony (figure 7A, Supplementary Material, video 1.0 1000.mp4 and video 0.3 10000.mp4).

**Figure 7:**
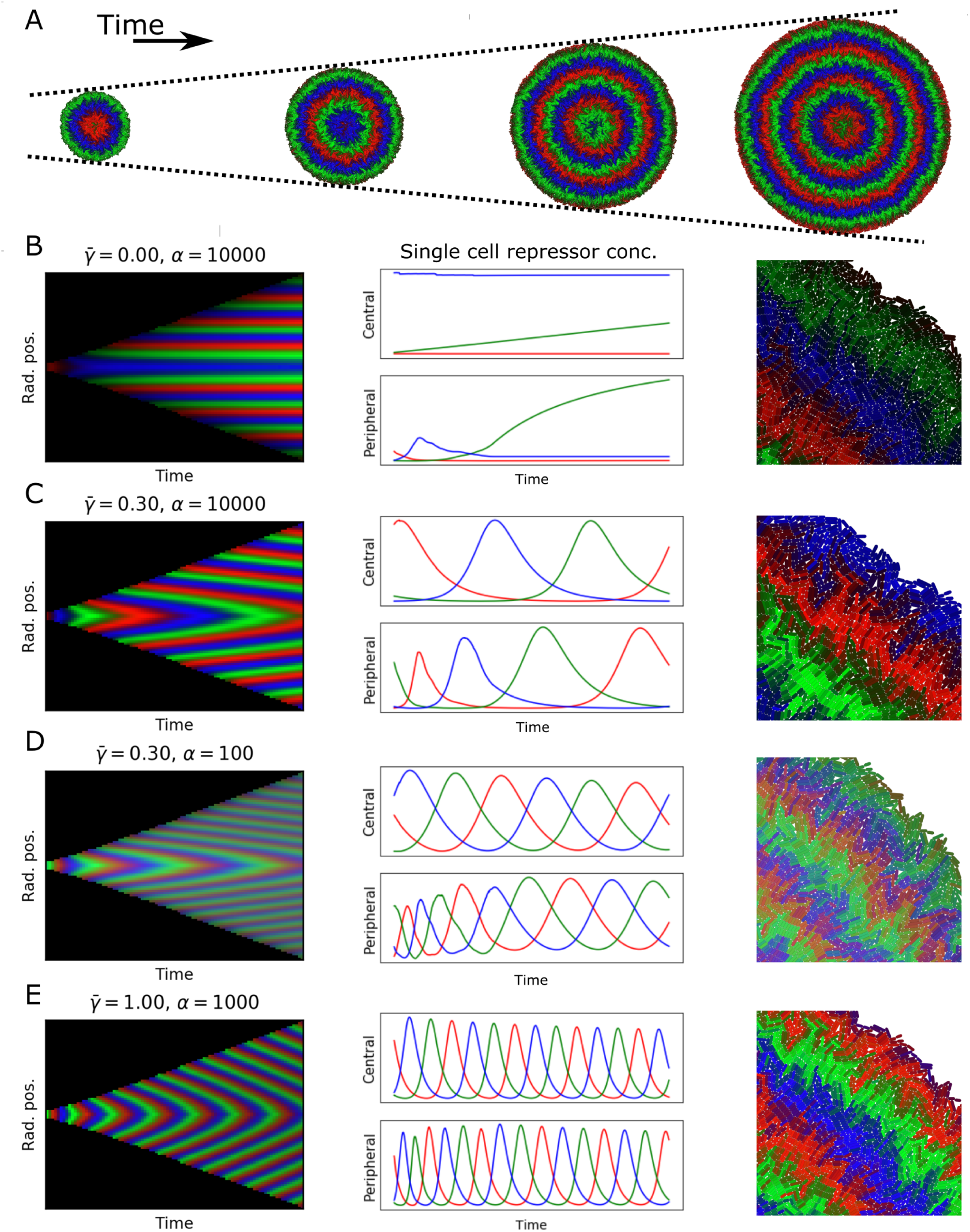
Simulations of growing colonies. **A**. Colonies with 5,000, 20,000, 35,000 and 50,000 cells, equally spaced 9.3 doubling times apart, with 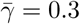, *α*=10.000. **(B-E)** Each panel shows: kymograph of repressor concentrations (51 doublings, approximately 60.000 cells); time dynamics for central cell and peripheral cell in colony; close up of edge of colony at end of experiment. Parameters: **B**. 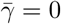, *α* = 10000. **(C)** 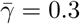, *α* = 10000. **D**. 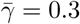, *α* = 100. **E**. 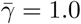, *α* = 1000.

In order to test the dependency of wavelength and wave speed on protein degradation and maximal expression rate, we simulated a range of 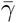 and *α* (kymographs in figure 7B-E). Our findings matched with the prediction of the 1D model; no waves were formed for 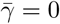, wave speed increased when 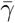 was increased, and wavelength increased when *α* was increased. The spatio-temporal dynamics of each repressor is regulated by protein dilution 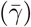, which moves the system from fixed rings (figure 7B) to an oscillatory behaviour which gives rise to traveling waves with different wavelengths controlled by maximum expression levels and speeds controlled by protein degradation (figure 7C-E).

Since we tracked every cell as they grow, replicate, and express proteins (figure 7B-E right column) we could follow the dynamics of individual cells as they move through the colony, changing their growth rate depending on their mechanical environment (figure 7B-E middle column). We selected central and peripheral cells for each of the colonies to study the most restricted and the most unconstrained cells. For colonies with travelling waves the constrained non-growing central cells oscillated with constant frequency and phase. Cells starting at the periphery of the colony however experience changes in growth rate as they move away from the colony edge that cause a sharp decrease in frequency and a resulting phase offset with respect to the central cell. We found that cells in the periphery exhibit higher frequency oscillations compared to central cells, and that difference is increased when increasing 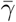 (figure 7 panels B-E central column). This is consistent with our theoretical prediction that growth rate reduction increases the period of oscillations, causing a phase offset between the interior and peripheral regions of the colony.

The wavelength and wave speed obtained from growing microcolonies was closely correlated to the predictions of our simple model (Pearson’s correlation coefficient 0.983 for wavelength and 0.999 for wave speed, figure 8). The length scale of wave speed and wavelength is set by *r*_0_, hence fitting a linear model between the predicted and simulated speed and wavelengths gives an estimate of *r*_0_ for the growing colony. From the wave speed values we obtained *r*_0_ = 11.6 and from wavelength *r*_0_ = 9.01, which is in close agreement with that estimated from the growth rate distribution of colonies (figure 2). Our one dimensional model predicts that the front velocity of the colony should be *v*_*front*_ = *r*_0_*/*2. From wave speed we obtain a value of *v*_*front*_ = 5.80 and from wavelength *v*_*front*_ = 4.51, which is extremely close to the estimated value of *v*_*front*_ = 5.00 for our individual based model (figure 2).

**Figure 8:**
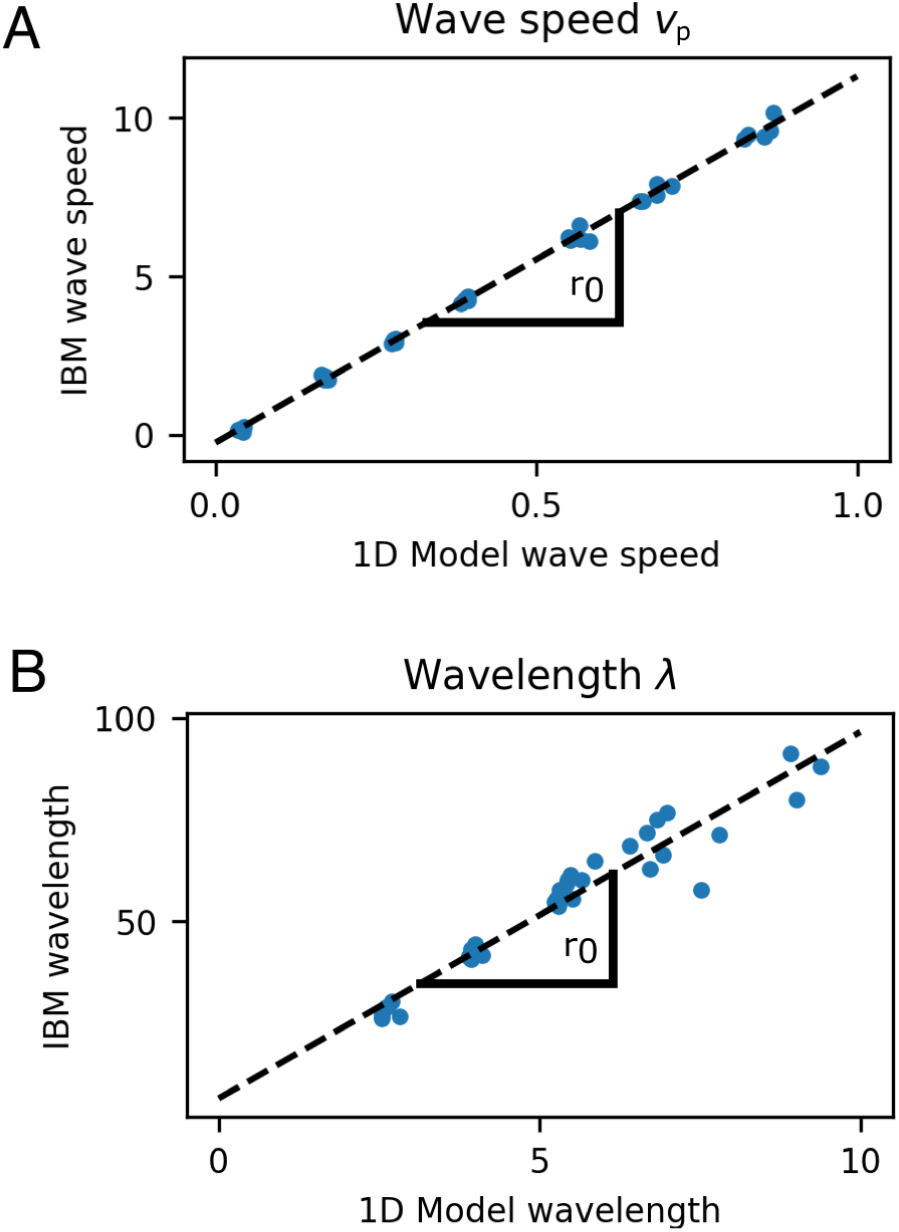
Comparison of wave speed (A) and wavelength (B) between 1D model and growing cell colonies (IBM). The length scale of wave speed and wavelength is set by *r*_0_ and the slope of these plots gives estimates for *r*_0_ as 11.6 (wave speed) and 9.01 (wavelength) cell diameters. Speed and wavelength of traveling waves were closely correlated between the models, with Pearson’s correlation coefficient 0.999 for wave speed, and 0.983 for wavelength.

### 2.7 Manipulating mechanical growth constraints to control pattern formation

Microfabricated cell culture devices and microfluidics provide fine control over the mechanical as well as biochemical conditions in which cells grow. As well as providing fresh nutrients via flow, maintaining cells in steady state, these devices provide techniques to physically constrain cell growth and therefore another mode of design of spatio-temporal patterns induced by growth rate heterogeneity as we have demonstrated. Commonly microfluidic devices are designed to constrain cells to a monolayer, while allowing loading of seed cells into a chamber or channel (figure 9A, B). We imposed two such constraints on our colonies. In a long thin channel (400 × 20 × 1 cell diameters) cells form one dimensional travelling waves directed along the channel axis (figure 9A). When constrained to a chamber (80 × 80 × 1 cell diameters) we observed the emergence of travelling waves during unconstrained growth (figure 9B, *t* + 4). These waves were sustained over long time periods after the cells became constrained and stopped growing altogether (figure 9B, *t* + 8, *t* + 12). Growth is necessary to form travelling waves but the established phase offsets between different radial positions are maintained after growth arrest, continuing to produce travelling waves. The history of the shape of wavefront is therefore retained in the pattern. Finally, we demonstrate that growth rate heterogeneity in three dimensional colonies also generates traveling waves as layers (figure 9C), showing that the spatio-temporal pattern is not specific to monolayers (see Supplementary video video 3d.mp4 and video 3d2.mp4).

**Figure 9:**
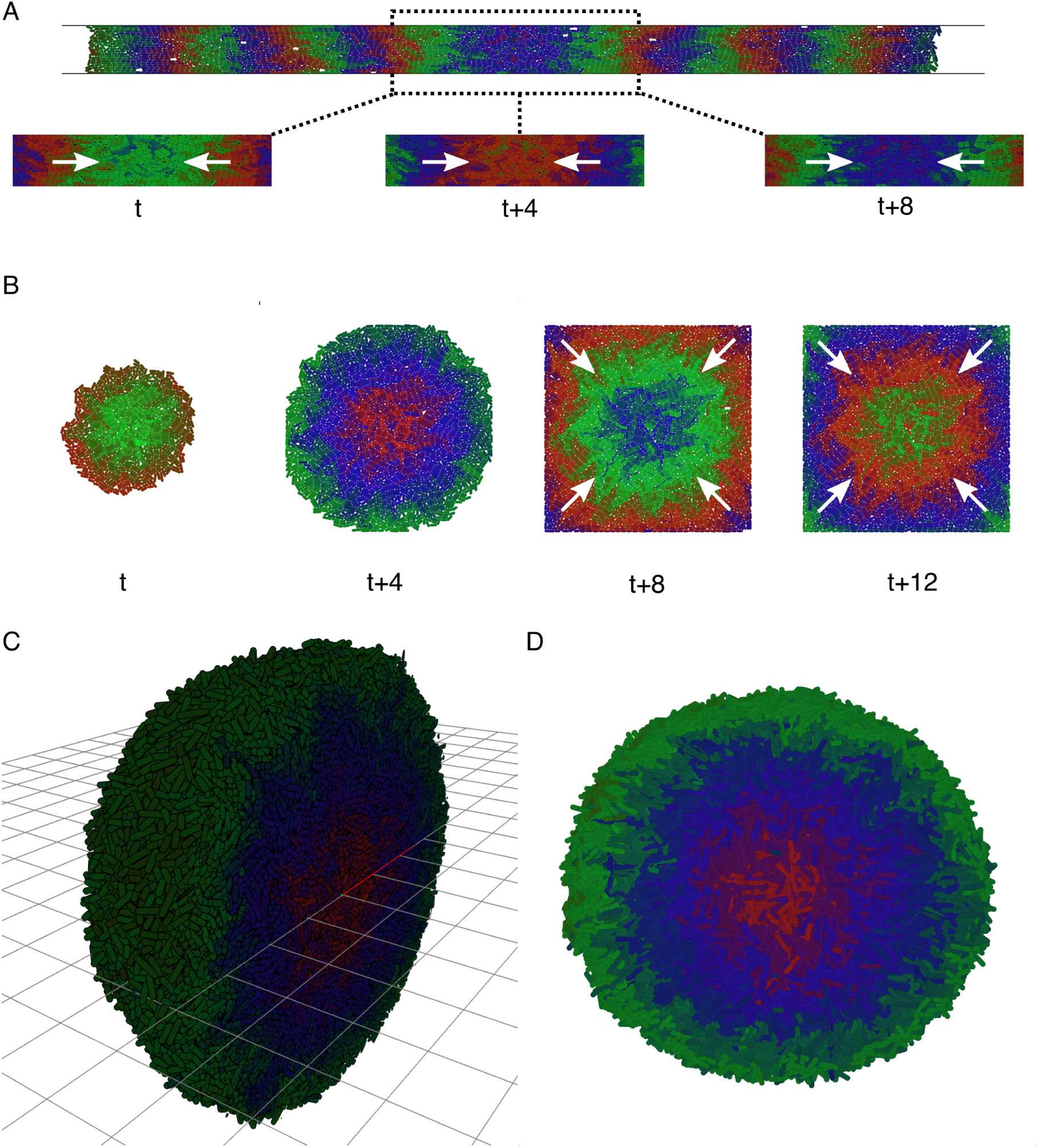
Traveling waves in microfluidic devices. **(A)**. Simulations of a monolayer of cells in a narrow infinite channel 20 cell diameters wide and one cell diameter tall. Traveling waves start at the sides and merge in the center (white arrows). **(B)**. Growing colony in a square simulated microfluidic chamber. Before the colony reaches the constraints it grows with radial symmetry and initiates travelling waves (t, t+4). At t+12 the colony has used all the space and is constrained by chamber, growth arrest occurs, but the travelling waves continue (white arrows). **(C)**. Spherical colonies grow when not constrained to a plane, forming travelling layers of gene expression. Image shows a cutaway of half the colony. **(D)**. Transversal section of C.

## 3 Discussion

Here we demonstrated how biophysical constraints on growth can induce spatio-temporal pattern formation from a simple genetic circuit. By coupling the repressilator [14] to a heterogeneous growth rate pattern via protein dilution we generated emergent travelling waves of gene expression. These travelling waves can be characterized by two properties; the wavelength and the wave speed. These properties are determined by two simple parameters that are feasible to control experimentally; the protein degradation rate, which controls the wave speed, and the maximal protein expression rate, which controls the wavelength. Our results make quantitative and qualitative predictions about the spatio-temporal patterns produced by the repressilator in growing cell colonies.

Our analysis predicts that travelling waves will be observed if the ratio of protein degradation to growth rate 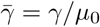, is sufficiently high for the waves to form in the time of the experiment. For 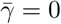 we predict the formation of static rings of gene expression as observed in experimenmts [14], however we show here that this pattern could be generated purely by protein dilution and does not require cells to transition into stationary phase. We show that increasing 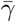, which means increasing protein degradation rate or decreasing growth rate, will increase the speed of the waves. This could be achieved by choosing one of several protein degradation tag sequences that target the proteins for proteolysis [49]. Further, our model suggests that increasing maximum protein expression rate *α*, for example by choosing a more efficient ribosome binding site (RBS) [51] will increase the wavelength of the pattern. We parameterize simple empirical models for the effects of each of these genetic design modifications; log10(*α*) = *λ* − 1 and 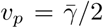. Thus we have effectively generated a quantitative datasheet for the repressilator gene circuit topology operating in a simple microcolony chassis.

We derive a simple model of coupling genetic circuits to growth rate via protein dilution, and show that it accurately predicts travelling wave patterns in growing cell colonies, their speed and wavelength. The model also accurately predicts the colony front velocity. The mathematical and computational approach outlined here is not specific to the repressilator nor to bacterial colonies and could make predictions about spatial patterns produced by other circuit topologies and chassis. Here we did not consider gene circuits that affect growth rate, for example by regulation of metabolism, which may produce more complex spatial patterns [22], however it could be included in our framework leading to a more complex set of coupled differential equations. Thus, this analysis approach implements the rational design of spatio-temporal patterns of gene expression, enabling the design stage of the design-build-test-learn cycle.

Oscillators are important in regulation of multicellular systems and many studies have reproduced oscillations in synthetic genetic circuits by assembling different devices combining modular parts [52, 53, 54, 55]. Studying and engineering synthetic oscillators can direct us to understand complex multicellular behaviours at multiple scales, in particular here the emergence of travelling waves of gene expression in populations of cells. There are a wide range of phenomena in which a key element to a developmental process is the appearance of a travelling wave of chemical concentration, mechanical deformation [56], electrical or other type of signal [57]. Two examples are the chemical and mechanical waves which propagate on the surface of many vertebrate eggs [58]. A developing embryo presents a large number of wave like events that appear after fertilisation[59]. Thus, one importance of this work is that we were able to rationally design and manipulate *in silico* genetic circuits to recapitulate such patterns with tunable wavelength and wave speed.

Noise is known to affect oscillators in various ways including stochastic coherence which makes the oscillations more consistent [60], and may therefore stabilize spatio-temporal patterns to stochastic fluctuations in gene expression. We do not consider the role of noise in this study because at the parameter values we explore, the stochastic behaviour approximates the continuous model, with regular and sustained synchronous oscillations (Supplementary Materials, figure S2). We also note that since [14] observed synchronous long-term oscillations that form ring patterns in growing colonies, cells must be synchronized on average over long length and time scales, and so noise is not likely to be important in the pattern formation process described here. However, it would be interesting to consider the role of noise in the generation of spatio-temporal patterns [61, 62] due to lower gene copy or other circuit properties.

We reason that the travelling waves described here are generated by phase and frequency changes induced by reduction in growth rate as cells become more distant from the edge of the colony, but maintained by protein degradation. In the absence of protein degradation no travelling waves but simple static rings form [14]. The phase differences are locked in as growth rate decays to zero, such that even after total growth arrest the travelling waves continue (figure 9B). The scale of the waves, their speed and their wavelength are determined by *r*_0_, the characteristic length scale of decay in growth rate. However, the radius of the colony also scales with *r*_0_ and so the overall pattern is in a sense scale invariant, showing the same relative wavelength and speed for any exponential growth rate profile.

Our results show that the speed of travelling waves in growing bacterial colonies is approximately 10 cell diameters per doubling time (approximately 10*µm* per hour for *E. coli*) towards the colony center, but the colony border grows at only around 4 cell diameters per doubling. Hence gene expression information can be transferred faster via a travelling wave than by the physical transmission of cells. The ability to tune the wavelength *λ* and wave speed *v*_*p*_ of these patterns could enable design of novel cell-cell communication systems based on oscillatory signals. Further, coupling the oscillator to production of pulses of diffusing chemicals such as acyl-homoserine lactones (AHLs) could be used to enhance information transmission [63, 64]. We note also that in a sense the travelling wave pattern, its speed and wavelength encode the history of the shape of the wavefront as the colony expands, which may be useful for example for information storage.

A fundamental result of this work is to demonstrate that mechanical constraint gives rise to higher order gene expression patterns in cell colonies, and provide such a system for analysis. There are a vast number of experimental conditions which could be created to induce different spatio-temporal patterns in such microcolonies. Microfluidics has shown to be of particular help to control mechanical constraints [65], substrate stiffness [66], nutrients [67], chemical inducers [13], cell-cell signaling [67] and pattern formation [68]. As we showed in figure 9, controlling biophysical constraints using different channel layouts and mechanical properties of the substrate could produce different patterns of growth rate that give rise to structures that mimic different stages of the development of organisms [25, 69]. A simple example is the one dimensional channel (figure 9A) which mimics in a simple way an embryo growing along its axis and sending back waves of gene expression from the front of the cell population.

In summary we report here novel travelling wave spatio-temporal patterns resulting from the growth rate dependent dynamics of a repressilator genetic oscillator circuit. We developed an analytical framework to predict the spatio-temporal behaviour of such genetic circuits in growing colonies. This framework allows multi-scale rational design of spatio-temporal patterns from genetic circuits and makes testable predictions for the synthetic biology design-build-test-learn cycle.

## 4 Material and Methods

Computation and analysis in this work were performed in Python [70] with the use of the packages NumPy [71], SciPy [72], Pandas [73], Jupyter [74], Matplotlib [75], Seaborn [76] and NetworkX [77].

### 4.1 Individual based model

We grew colonies from 1 up to 60,000 cells, simulated using CellModeller [32] with parameters Γ = 10 and Δ*t* = 0.05. Γ = *γ*_*cell*_*/γ*_*s*_ is the ratio of cell stiffness to substrate stiffness, which we estimate order of magnitude 10 (see Supplementary Material). Briefly, CellModeller simulates cells as rods extending along their axis but otherwise rigid. As cells expand the resulting constraint energy is minimized to find the new arrangement of cells. Cells experience viscous drag with the substrate (*γ*_*s*_) and along their length axis (*γ*_*cell*_), and divide when they reach a target length set to *l*_0_ = 3.5 cell diameters. At division the cell is divided into two equal sized rods, which are randomly perturbed slightly in their axis orientation. Cells were constrained to lie in a plane, except in figure 9C in which cells grew in three dimensions.

Colonies were grown for approximately 48 doubling times and the final radius of the colonies was approximately ∼ 230 cell diameters. Since these simulations corresponds to *E. Coli* cells, these units represents approximately ∼ 230*µm*. Simulations were stored in a file for each time step. This file contains information about the state of each cell present in the colony, including the position, protein concentrations, growth rate, volume, length, among other variables.

### 4.2 Colony growth analysis

Growth rate *µ* and radial position *R* were obtained for each cell from 3 growing colonies from 5.000 up to 60,000 cells. At each time point the colony radius *R*_*max*_(*t*) was calculated and divided into *n* bins of size Δ*r* = 5 cell diameters from the edge *b*_0_ to the center *b*_*n*_. The growth rates *µ* of all cells with *r* ∈ [*b*_*n*_, *b*_*n*+1_) were averaged to get ⟨*µ*⟩. An exponential of the form *e*^−*r*(*t*)*k*^ was fitted to ⟨*µ*⟩ at each time, obtaining *k* at colonies with different *R*_*max*_. A linear model was fitted to the colony radius *R*_*max*_(*t*) when *t* > 10 to compute the front velocity *v*_*front*_ using numpy.polyfit.

### 4.3 Kymograph construction for 1D model

To obtain values of *p*_*i*_(*r, t*) we integrate equation 11. Starting from some initial colony radius *R*_0_ we initialize *p*_*i*_(*r*, 0) for *r* a regularly spaced lattice on (0, *R*_0_). We use *p*_2_(*r*, 0) = 5 for all *r*, that is homogeneous expression of only *p*_2_. At each step of an explicit Euler integration scheme we find the new cell positions and construct a new regularly spaced lattice in (0, *R*(*t*)) = (0, *R*_0_ + *t/*2) and interpolate *p*_*i*_(*r, t* + Δ*t*) onto this grid before taking the next integration step. The algorithm is as follows:

1. Initialize *r*(0) as a regular lattice on (0, *R*_0_) and 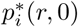 to some values.
2. Compute *p*_*i*_(*r, t* + Δ*t*) by an explicit Euler step of equation 11, and *r*(*t* + Δ*t*) using equation 6.
3. Compute a new regular mesh *r*′(*t* + Δ*t*) on (0, *R*(*t* + Δ*t*)).
4. Interpolate the protein concentrations to get *p*_*i*_(*r*′, *t* + Δ*t*).
5. Set *t* → *t* + Δ*t*, and *r*(*t* + Δ*t*) → *r*′(*t* + Δ*t*), and repeat from step 2.

At the end of this procedure we have constructed a set of samples *p*_*i*_(*r, t*) which we then interpolate to form the kymograph.

### 4.4 Dynamical simulations of gene expression

Using the file stored for each simulation in the IBM, we have a representation of the biophysical model decoupled from the genetic circuit. Using these data we performed simulations of the gene expression model derived in equation 11. In order to keep updating the state of the cells, which is affected by cell division, we constructed a graph of parent-child relations. Thus, we integrate equation 11 forward using explicit Euler method between each state of the biophysical model. One assumption made is that when a cell divides the children inherit the value of the protein concentration his parent. We assume the number of proteins divides equally between the two cells, as does the volume of the cell, keeping protein concentration constant. Resultant simulations then serialized to a JSON file. These files were later used to perform analysis and create visual representations.

### 4.5 Kymograph construction for individual based simulations

Using the JSON file obtained in the temporal simulation of gene expression with the biophysical model, we generated positions and growth rates of cells. Then we binned cells according to their radial position using bin size Δ*R* = 5. Finally we take average protein concentration in each bin and repeat for all time steps to get *p*_*i*_(*R, t*).

### 4.6 Wave speed estimation

First we take each row of the kymograph and identify the radial peaks (scipy.signal.find peaks) in each protein concentration for each time step. Next the peaks are paired with the nearest peak in the previous time step, and the average distance between them used to calculate the wave speed as *v* = ⟨Δ *x*⟩ */δt*, where ⟨Δ*x*⟩ is the mean peak to peak distance and *δt* is the simulation time step.

### 4.7 Wavelength estimation

In order to estimate the wavelength *λ* of the travelling waves we note that *λ* = (*v* + *v*_*front*_)*T*, where *v* is the wave speed, *T* is the period of oscillations, and 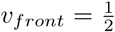 is the velocity of the colony edge or wavefront. To estimate the oscillation period we take the leading edge of the colony and compute the peaks in its time varying protein concentration *p*_*i*_(*r* = 0, *t*). Then as above we estimate the period as the mean of the peak to peak times so that *T* = ⟨Δ*t*⟩. The wave speed is taken from the calculations described above, and the resulting estimate for wavelength is 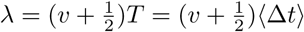.

### 4.8 Analytical estimates of wavelength and wave speed

We used equations 14-15 to estimate the wavelength and wave speed of traveling waves that the repressilator would produce with an exponential growth rate profile (equation 1). First we numerically integrated equation 11 with fixed growth rate 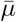. For each combination of parameters we simulated oscillations at the colony edge 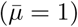 and the colony interior 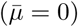, and measured the periods *T*_*ext*_ and *T*_*int*_ as described above. These values were then substituted into equations 14-15 to compute the estimated wavelength and wave speed.

## Author Contributions

GYF, GV and TJR designed the study and analyzed the data. MMS, GYF, GV and TJR wrote the manuscript.

## Acknowledgments

We thank Gustavo Düring, Luca Ciandrini, Pascal Rogalla, and Ignacio Medina for helpful and stimulating discussions. We also thank the members of the Synthetic Biology Lab for their support and encouragenent - Anibal Arce, Kevin Simpson, Tamara Matute, Isaac Nuñez, Fernán Federici, among others.

## Supplemental Data

Supplementary Material and videos are available separately.

## Data Availability Statement

The datasets generated and analyzed, as well as the code to perform the analysis and plots presented here are available at https://github.com/SynBioUC/SpatialOscillator.

## References

[1] Kyle H. Vining and David J. Mooney. “Mechanical forces direct stem cell behaviour in development and regeneration”. In: Nature Reviews Molecular Cell Biology 18.12 (2017), pp. 728–742. ISSN: 14710080. doi: 10.1038/nrm.2017.108. URL: http://dx.doi.org/10.1038/nrm.2017.108.

[2] Chii J. Chan, Carl Philipp Heisenberg, and Takashi Hiiragi. “Coordination of Morphogenesis and Cell-Fate Specification in Development”. In: Current Biology 27.18 (2017), R1024–R1035. ISSN: 09609822. DOI: 10.1016/j.cub.2017.07.010.

[3] Javier Santos-Moreno and Yolanda Schaerli. “Using Synthetic Biology to Engineer Spatial Patterns”. In: Advanced Biosystems 3.4 (2019), pp. 1–15. ISSN: 23667478. DOI: 10.1002/adbi.201800280.

[4] A Aziz Aboobaker et al. “Drosophila microRNAs exhibit diverse spatial expression patterns during embryonic development”. In: Proceedings of the National Academy of Sciences 102.50 (2005), pp. 18017–18022. DOI: 10.1073/pnas.0508823102.

[5] Evgeniy Khain and Leonard M Sander. “Dynamics and pattern formation in invasive tumor growth”. In: Physical review letters 96.18 (2006), p. 188103. DOI: 10.1103/PhysRevLett.96.188103.

[6] Margaret J Velardo et al. “Patterns of gene expression reveal a temporally orchestrated wound healing response in the injured spinal cord”. In: Journal of Neuroscience 24.39 (2004), pp. 8562–8576. DOI: 10.1523/JNEUROSCI.3316-04.2004.

[7] Nikolce Gjorevski and Celeste M Nelson. “Endogenous patterns of mechanical stress are required for branching morphogenesis”. In: Integrative Biology 2.9 (2010), pp. 424–434. DOI: 10.1039/c0ib00040j.

[8] Mingqi Xie and Martin Fussenegger. “Designing cell function: assembly of synthetic gene circuits for cell biology applications”. In: Nature Reviews Molecular Cell Biology 19.8 (2018), pp. 507–525. ISSN: 14710080. DOI: 10.1038/s41580-018-0024-z. URL: http://dx.doi.org/10.1038/s41580-018-0024-z.

[9] Timothy S Gardner, Charles R Cantor, and James J Collins. “Construction of a genetic toggle switch in *Escherichia coli*”. In: Nature 403.6767 (2000), pp. 339–342. DOI: 10.1038/35002131.

[10] Enoch Yeung et al. “Biophysical Constraints Arising from Compositional Context in Synthetic Gene Networks”. In: Cell Systems 5.1 (2017), 11–24.e12. ISSN: 24054720. DOI: 10.1016/j.cels.2017.06.001. URL: http://dx.doi.org/10.1016/j.cels.2017.06.001.

[11] Michael B Elowitz and Stanislas Leibler. “A synthetic oscillatory network of transcriptional regulators”. In: Nature 403.6767 (2000), pp. 335–338. DOI: 10.1038/35002125.

[12] Jesse Stricker et al. “A fast, robust and tunable synthetic gene oscillator”. In: Nature 456.7221 (2008), pp. 516–519. DOI: 10.1038/nature07389.

[13] Tal Danino et al. “A synchronized quorum of genetic clocks”. In: Nature 463.7279 (2010), pp. 326–330. DOI: 10.1038/nature08753.

[14] Laurent Potvin-Trottier et al. “Synchronous long-term oscillations in a synthetic gene circuit”. In: Nature 538.7626 (2016), pp. 514–517. DOI: 10.1038/nature19841.

[15] Alvin Tamsir, Jeffrey J Tabor, and Christopher A Voigt. “Robust multicellular computing using genetically encoded NOR gates and chemical ‘wires’”. In: Nature 469.7329 (2011), pp. 212–215. DOI: 10.1038/nature09565.

[16] Alec AK Nielsen et al. “Genetic circuit design automation”. In: Science 352.6281 (2016), aac7341. DOI: 10.1126/science.aac7341.

[17] Alexander A Green et al. “Complex cellular logic computation using ribocomputing devices”. In: Nature 548.7665 (2017), pp. 117–121. DOI: 10.1038/nature23271.

[18] Jongmin Kim, Peng Yin, and Alexander A. Green. “Ribocomputing: Cellular Logic Computation Using RNA Devices”. In: Biochemistry 57.6 (2018), pp. 883–885. ISSN: 15204995. DOI: 10.1021/acs.biochem.7b01072.

[19] Adison Wong et al. “Layering genetic circuits to build a single cell, bacterial half adder”. In: BMC biology 13.1 (2015), p. 40. DOI: 10.1186/s12915-015-0146-0.

[20] David Ausländer et al. “Programmable full-adder computations in communicating three-dimensional cell cultures”. In: Nature methods 15.1 (2018), p. 57. DOI: 10.1038/nmeth.4505.

[21] Nan Luo, Shangying Wang, and Lingchong You. “Synthetic Pattern Formation”. In: Biochemistry 58.11 (2019), pp. 1478–1483. ISSN: 15204995. DOI: 10.1021/acs.biochem.8b01242.

[22] Isaac N Nuñez et al. “Artificial symmetry-breaking for morphogenetic engineering bacterial colonies”. In: ACS synthetic biology 6.2 (2017), pp. 256–265. DOI: 10.1021/acssynbio.6b00149.

[23] David Karig et al. “Stochastic Turing patterns in a synthetic bacterial population”. In: Proceedings of the National Academy of Sciences 115.26 (2018), pp. 6572–6577. ISSN: 0027-8424. DOI: 10.1073/pnas.1720770115.

[24] Timothy J Rudge et al. “Cell polarity-driven instability generates self-organized, fractal patterning of cell layers”. In: ACS synthetic biology 2.12 (2013), pp. 705–714. DOI: 10.1021/sb400030p.

[25] Satoshi Toda et al. “Programming self-organizing multicellular structures with synthetic cell-cell signaling”. In: Science 361.6398 (2018), pp. 156–162. ISSN: 10959203. DOI: 10.1126/science.aat0271.

[26] C. P. Healy and T. L. Deans. “Genetic circuits to engineer tissues with alternative functions”. In: Journal of Biological Engineering 13.1 (2019), pp. 1–7. ISSN: 17541611. DOI: 10.1186/s13036-019-0170-7.

[27] Natalie A. Cookson et al. “Queueing up for enzymatic processing: Correlated signaling through coupled degradation”. In: Molecular Systems Biology 7.1 (2011). ISSN: 17444292. DOI: 10.1038/msb.2011.94.

[28] Paul J. Steiner et al. “Criticality and Adaptivity in Enzymatic Networks”. In: Biophysical Journal 111.5 (2016), pp. 1078–1087. ISSN: 15420086. DOI: 10.1016/j.bpj.2016.07.036. URL: http://dx.doi.org/10.1016/j.bpj.2016.07.036.

[29] Drew Endy and Roger Brent. “Modelling cellular behaviour”. In: Nature 409.6818 (2001), pp. 391–395. DOI: 10.1038/35053181.

[30] Hidde De Jong. “Modeling and simulation of genetic regulatory systems: A literature review”. In: Journal of Computational Biology 9.1 (2002), pp. 67–103. ISSN: 10665277. DOI: 10.1089/10665270252833208.

[31] Thomas E Gorochowski. “Agent-based modelling in synthetic biology”. In: Essays in biochemistry 60.4 (2016), pp. 325–336. DOI: 10.1042/EBC20160037.

[32] Timothy J. Rudge et al. “Computational Modeling of Synthetic Microbial Biofilms”. In: ACS Synthetic Biology 1.8 (Aug. 2012), pp. 345–352. ISSN: 2161-5063. DOI: 10.1021/sb300031n. URL: http://pubs.acs.org/doi/10.1021/sb300031n.

[33] J J Sickle et al. “Integrative Circuit-Host Modeling of a Genetic Switch in Varying Environments”. In: May (2020), pp. 1–13. DOI: 10.1038/s41598-020-64921-5.

[34] FDC Farrell et al. “Mechanically driven growth of quasi-two-dimensional microbial colonies”. In: Physical review letters 111.16 (2013), pp. 1–5.

[35] Matthew AA Grant et al. “The role of mechanical forces in the planar-to-bulk transition in growing *Escherichia coli* microcolonies”. In: Journal of The Royal Society Interface 11.97 (2014), p. 20140400. DOI: 10.1098/rsif.2014.0400.

[36] Tamás Vicsek, Miklós Cserző, and Viktor K Horváth. “Self-affine growth of bacterial colonies”. In: Physica A: Statistical Mechanics and its Applications 167.2 (1990), pp. 315–321. DOI: 10.1016/0378-4371(90)90116-A.

[37] William PJ Smith et al. “Cell morphology drives spatial patterning in microbial communities”. In: Proceedings of the National Academy of Sciences 114.3 (2017), E280–E286. DOI: 10.1073/pnas.1613007114.

[38] Peter Neubauer, HY Lin, and B Mathiszik. “Metabolic load of recombinant protein production: inhibition of cellular capacities for glucose uptake and respiration after induction of a heterologous gene in *Escherichia coli*”. In: Biotechnology and bioengineering 83.1 (2003), pp. 53–64. DOI: 10.1002/bit.10645.

[39] Stefan Klumpp, Zhongge Zhang, and Terence Hwa. “Growth rate-dependent global effects on gene expression in bacteria”. In: Cell 139.7 (2009), pp. 1366–1375. DOI: 10.1016/j.cell.2009.12.001.

[40] Stefan Klumpp. “Growth-rate dependence reveals design principles of plasmid copy number control”. In: PloS one 6.5 (2011). DOI: 10.1371/journal.pone.0020403.

[41] Klaus B Andersen and Kaspar von Meyenburg. “Are growth rates of *Escherichia coli* in batch cultures limited by respiration?” In: Journal of bacteriology 144.1 (1980), pp. 114–123. DOI: 6998942.

[42] Mitsugu Matsushita and Hiroshi Fujikawa. “Diffusion-limited growth in bacterial colony formation”. In: Physica A: Statistical Mechanics and its Applications 168.1 (1990), pp. 498–506. DOI: 10.1016/0378-4371(90)90402-E.

[43] Hannah H. Tuson et al. “Measuring the stiffness of bacterial cells from growth rates in hydrogels of tunable elasticity”. In: Molecular Microbiology 84.5 (2012), pp. 874–891. ISSN: 0950382X. DOI: 10.1111/j.1365-2958.2012.08063.x.

[44] James J. Winkle et al. “Modeling mechanical interactions in growing populations of rod-shaped bacteria”. In: Physical Biology 14.5 (2018). ISSN: 14783975. DOI: 10.1088/1478-3975/aa7bae. eprint: 1702.08091.

[45] JV Matson and William G Characklis. “Diffusion into microbial aggregates”. In: Water Research 10.10 (1976), pp. 877–885. DOI: 10.1016/0043-1354(76)90022-1.

[46] Steven P Fraleigh and Henry R Bungay. “Modelling of nutrient gradients in a bacterial colony”. In: Microbiology 132.7 (1986), pp. 2057–2060. DOI: 10.1099/00221287-132-7-2057.

[47] Thomas Guélon, Jean-Denis Mathias, and Guillaume Deffuant. “Influence of spatial structure on effective nutrient diffusion in bacterial biofilms”. In: Journal of biological physics 38.4 (2012), pp. 573–588. DOI: 10.1007/s10867-012-9272-x.

[48] Zhicheng Long et al. “Microfluidic chemostat for measuring single cell dynamics in bacteria”. In: Lab on a Chip 13.5 (2013), pp. 947–954. DOI: 10.1039/c2lc41196b.

[49] Oliver Purcell et al. “Temperature dependence of ssrA-tag mediated protein degradation”. In: Journal of biological engineering 6.1 (2012), p. 10. DOI: 10.1186/1754-1611-6-10.

[50] Matteo Osella and Marco Cosentino Lagomarsino. “Growth-rate-dependent dynamics of a bacterial genetic oscillator”. In: Physical Review E 87.1 (2013), p. 012726. DOI: 10.1103/PhysRevE.87.012726.

[51] Howard M. Salis, Ethan A. Mirsky, and Christopher A. Voigt. “Automated design of synthetic ribosome binding sites to control protein expression”. In: Nature Biotechnology 27.10 (2009), pp. 946–950. ISSN: 10870156. DOI: 10.1038/nbt.1568.

[52] Henrike Niederholtmeyer et al. “Rapid cell-free forward engineering of novel genetic ring oscillators”. In: eLife 4.OCTOBER2015 (2015), pp. 1–18. ISSN: 2050084X. DOI: 10.7554/eLife.09771.

[53] Jintao Liu et al. “Metabolic co-dependence gives rise to collective oscillations within biofilms”. In: Nature 523.7562 (2015), pp. 550–554. ISSN: 14764687. DOI: 10.1038/nature14660.

[54] Ruben Perez-Carrasco et al. “Combining a Toggle Switch and a Repressilator within the AC-DC Circuit Generates Distinct Dynamical Behaviors”. In: Cell Systems 6.4 (2018), 521–530.e3. ISSN: 24054720. DOI: 10.1016/j.cels.2018.02.008. URL: https://doi.org/10.1016/j.cels.2018.02.008.

[55] David T. Riglar et al. “Bacterial variability in the mammalian gut captured by a single-cell synthetic oscillator”. In: Nature Communications 10.1 (2019), pp. 1–12. ISSN: 20411723. DOI: 10.1038/s41467-019-12638-z. URL: http://dx.doi.org/10.1038/s41467-019-12638-z.

[56] David R. Espeso et al. “Physical forces shape group identity of swimming *Pseudomonas putida* cells”. In: Frontiers in Microbiology 7.SEP (2016), pp. 1–11. ISSN: 1664302X. DOI: 10.3389/fmicb.2016.01437.

[57] James D. Murray. Mathematical Biology I. An Introduction. 3rd ed. Vol. 17. Interdisciplinary Applied Mathematics. New York: Springer, 2002. DOI: 10.1007/b98868.

[58] Victoria E. Deneke and Stefano Di Talia. “Chemical waves in cell and developmental biology”. In: Journal of Cell Biology 217.4 (2018), pp. 1193–1204. ISSN: 15408140. DOI: 10.1083/jcb.201701158.

[59] Charles B. Kimmel et al. “Stages of embryonic development of the zebrafish”. In: Developmental Dynamics 203.3 (1995), pp. 253–310. ISSN: 10970177. DOI: 10.1002/aja.1002030302.

[60] Robert C Hilborn and Jessie D Erwin. “Stochastic coherence in an oscillatory gene circuit model”. In: Journal of theoretical biology 253.2 (2008), pp. 349–354.

[61] Francesc Saguès, José M. Sancho, and Jordi García-Ojalvo. “Spatiotemporal order out of noise”. In: Reviews of Modern Physics 79.3 (2007), pp. 829–882. ISSN: 00346861. DOI: 10.1103/RevModPhys.79.829.

[62] Tianshou Zhou et al. “Synchronization of genetic oscillators”. In: Chaos 18.3 (2008). ISSN: 10541500. DOI: 10.1063/1.2978183.

[63] John J Hopfield. “Kinetic proofreading: a new mechanism for reducing errors in biosynthetic processes requiring high specificity”. In: Proceedings of the National Academy of Sciences 71.10 (1974), pp. 4135–4139. DOI: 10.1073/pnas.71.10.4135.

[64] Shmoolik Mangan, Alon Zaslaver, and Uri Alon. “The coherent feedforward loop serves as a sign-sensitive delay element in transcription networks”. In: Journal of molecular biology 334.2 (2003), pp. 197–204. DOI: 10.1016/j.jmb.2003.09.049.

[65] Verena Ruprecht et al. “How cells respond to environmental cues - insights from bio-functionalized substrates”. In: Journal of Cell Science 130.1 (2017), pp. 51–61. ISSN: 14779137. DOI: 10.1242/jcs.196162.

[66] Weixing Wang et al. “A Microfluidic Hydrogel Chip with Orthogonal Dual Gradients of Matrix Stiffness and Oxygen for Cytotoxicity Test”. In: Biochip Journal 12.2 (2018), pp. 93–101. ISSN: 20927843. DOI: 10.1007/s13206-017-2202-z.

[67] Razan N. Alnahhas et al. “Spatiotemporal Dynamics of Synthetic Microbial Consortia in Microfluidic Devices”. In: ACS Synthetic Biology 8.9 (2019), pp. 2051–2058. ISSN: 21615063. DOI: 10.1021/acssynbio.9b00146.

[68] Vasily Kantsler et al. “Pattern engineering of living bacterial colonies using meniscus-driven fluidic channels”. In: (2020). DOI: 10.1021/acssynbio.0c00146. eprint: 1910.01595. URL: http://arxiv.org/abs/1910.01595.

[69] Marion B. Johnson, Alexander R. March, and Leonardo Morsut. “Engineering multicellular systems: Using synthetic biology to control tissue self-organization”. In: Current Opinion in Biomedical Engineering 4.October (2017), pp. 163–173. ISSN: 24684511. DOI: 10.1016/j.cobme.2017.10.008. URL: https://doi.org/10.1016/j.cobme.2017.10.008.

[70] Guido Van Rossum and Fred L. Drake. Python 3 Reference Manual. Scotts Valley, CA: CreateSpace, 2009. ISBN: 1441412697.

[71] Travis E Oliphant. A guide to NumPy. Vol. 1. Trelgol Publishing USA, 2006.

[72] Pauli Virtanen et al. “SciPy 1.0: fundamental algorithms for scientific computing in Python”. In: Nature methods 17 (2020). DOI: 10.1038/s41592-019-0686-2.

[73] Wes McKinney. “Data Structures for Statistical Computing in Python”. In: Proceedings of the 9th Python in Science Conference. Ed. by Stéfan van der Walt and Jarrod Millman. 2010, pp. 56–61. DOI: 10.25080/Majora-92bf1922-00a.

[74] Fernando Pérez and Brian E. Granger. “IPython: a System for Interactive Scientific Computing”. In: Computing in Science and Engineering 9.3 (May 2007), pp. 21–29. ISSN: 1521-9615. DOI: 10.1109/MCSE.2007.53. URL: https://ipython.org.

[75] J. D. Hunter. “Matplotlib: A 2D graphics environment”. In: Computing in Science & Engineering 9.3 (2007), pp. 90–95. DOI: 10.1109/MCSE.2007.55.

[76] Michael Waskom et al. “Seaborn: v0.8.1”. Version v0.8.1. In: (Sept. 2017). DOI: 10.5281/zenodo.883859. URL: https://doi.org/10.5281/zenodo.883859.

[77] Aric A. Hagberg, Daniel A. Schult, and Pieter J. Swart. “Exploring network structure, dynamics, and function using NetworkX”. In: Proceedings of the 7th Python in Science Conference. Ed. by Travis Vaught Gäel Varoquaux and Jarrod Millman. 2018, pp. 11–15.

