## Supplementary methods for "Novel tunable spatio-temporal patterns from a simple genetic oscillator circuit"

July 6, 2020

#### 1 Parameter estimates

- Cell volume ($V$) varies with growth rate, for a cell with $1\mu m$ height, $1\mu m$ width and $2\mu m$ length the cell volume is $2\mu m^3$, we consider a cell volume of $1\mu m^3$ or $10^{-15}$ litres for simplicity which is in the same order of magnitude churchward1981growth
- Constitutive transcription ($a$), 0.5 transcripts per second elowitz2000synthetic. Which is in the range of 0.1 to 1 transcripts per second liang1999activities We considered 1 transcript as a particle so 0.5 particles are $8.3 \times 10^{-25}$ moles, divided by cell volume to obtain $8.3 \times 10^{-10}$
- Leaky transcription rate ($b$), $5 \times 10^{-4}$ transcripts per second elowitz2000synthetic. We considered 1 transcript as a particle so $5 \times 10^{-4}$ are $8.30 \times 10^{-28}$ moles, divided by cell volume to obtain a concentration rate of $8.30 \times 10^{-13}$ Molar/second.
- The average length of repressors LacI, TetR and $\lambda$ cl is 871 bp elowitz2000synthetic, 290.3 codons or amino acids when translated, similar to dsRed monomer with 220 amino acids. Translation rate ($c$), 6.7 DsRed monomers per transcript per minute guet2008minimally. We consider proteins and transcripts as particles, we get particle over particle per 60s, particles get cancelled and we get a translation rate of $1.11 \times 10^{-1} s^{-1}$.
- The switching concentration ($K$), 40 repressors per cell elowitz2000synthetic We consider proteins as a particle so 40 repressor protein are $6.64 \times 10^{-23}$ moles, divided by cell volume to obtain a concentration of $6.64 \times 10^{-8} Molar$.
- Growth rate ranges from $\mu = 0.2h^{-1}$ to $1.2h^{-1}$ andersen1980growth. We consider $\mu = 1h^{-1}$ or $2.77 \times 10^{-4} s^{-1}$.
- mRNA half life is estimated as 5-10 minutes taniguchi2010quantifying giving a degradation rate of $\delta = log(2)/5 = 0.139$ per minute, which is $2.31 \times 10^{-3} s^{-1}$.
- Protein half life is taken from purcell2012temperature as 41 minutes, giving degradation rate $log(2)/41 = 0.017$ which is $2.8 \times 10^{-4}$.
- The Young's modulus of *E. coli* cells at short time scales, which in effect relates to the effective viscosity at long time scales of growth, has been estimated between $10^5$ and $10^8$ Pa. Typical hydrogels and PDMS achieve stifness in the range $10^3$ Pa to $10^6$ MPa seghir2015extended. (seghi just show range between 0.8Mpa and 10Mpa) The biophysical gamma, lets call it $\Gamma$ is the ratio of cell to substrate stiffness. If we take PDMS stiffness of $10^6$ Pa and cell stiffness then $\Gamma$ is in the range $10^{-1}$ and $10^2$. Taking typical values we could justify using 0.1, 1, and 10.

| Parameter | Meaning | Order of magnitude |
|---|---|---|
| a | Constitutive transcription rate of repressor genes | $10^{-9} Molar\ s^{-1} [c1]$ |
| b | Leaky transcription rate of repressor genes | $10^{-13} Molar\ s^{-1} [c2]$ |
| c | Translation rate of repressor genes | $10^{-1} s^{-1} [c2, c3]$ |
| K | Switching concentration of repressors | $10^{-7} Molar [c2]$ |
| n | Cooperativity of repressors | $10^{0}$ [c2] |
| $\delta$ | Degradation rate of mRNA of repressor | $10^{-3} s^{-1} [c2]$ |
| $\gamma$ | Degradation rate of repressor | $10^{-4} s^{-1} [c2]$ |
| $\mu$ | Growth rate of *Escherichia coli* | $10^{-4} s^{-1} [c4]$ |

Table 1: Parameters of the two-step model of the repressilator. [c1]= churchward1981growth, [c2]= elowitz2000synthetic, [c3]= guet2008minimally, [c4]= andersen1980growth.

### 2 Estimated value of $\alpha$ and $\bar{\gamma}$

Using the order of magnitude estimates in table 1 we have,

$$\alpha = \frac{ac}{\delta \mu_0 K} \approx \frac{10^{-9} 10^{-1}}{10^{-3} 10^{-4} 10^{-7}} = 10^4 \tag{1}$$

For $\bar{\gamma}$ we have simply $\bar{\gamma} = \gamma/\mu_0 \approx 10^{-4}/10^{-4} = 1$. Based on these estimates we chose $\alpha \in (10^2, 10^5)$ and $\bar{\gamma} \in (10^{-3}, 1)$. To construct heatmaps of wavelength and wave speed we computed solutions for $25 \times 25$ uniformly log spaced values of $\bar{\gamma}$ and $\alpha$.

### 3 Growth rate profile

We simulated growing colonies using accurate individual base model, CellModeller (CM) Rudge2012 with $\Gamma = 10$ and $\Delta t = 0.05$, to extract growth rate given the biophysical constraints. We sample colonies from 5,000 cells with steps of 5,000 until 60,000 cells. Each colony size is characterized by its radius $R_{max}$. In 1A, we can see the characteristic exponential decrease on growth rate in colonies of different sizes. In 1B, we can see how exponential decrease shape is conserved across the different colony sizes when compared in terms of distance from edge $r$.

### 4 Stochastic simulation

We simulated the model of [1] which consists of a simple one step protein production and degradation, and plasmid replication and degradation, using the Gillespie stochastic simulation algorithm gillespie1977exact. The parameters of the model are $K$ the switching point in number of repressor proteins per cell, $\lambda$ the maximum protein expression rate, $n$ the cooperativity, $b$ the mean protein burst size, $N_0$ the steady state number of plasmids. We set parameters $K = 10^3$, $\lambda = 10^5$, $n = 2$, $b = 1$, $N_0 = 100$. These parameters correspond to $\alpha = 10^4$ and $\bar{\gamma} + \bar{\mu} = 1$ for our continuous model, which we showed are physiologically reasonable. We simulated 100 independent trajectories of this stochastic model showing that they sustain synchronized oscillations over at least 50 generations (figure 2).

### 5 Kymographs of individual based model

Colonies were grown from 1 to approximately 60,000 cells and radial averaging used to compute the kymograph, representing the spatio-temporal dynamics of the system, for a range of parameters $\bar{\gamma}$ and $\alpha$ (figure 3).

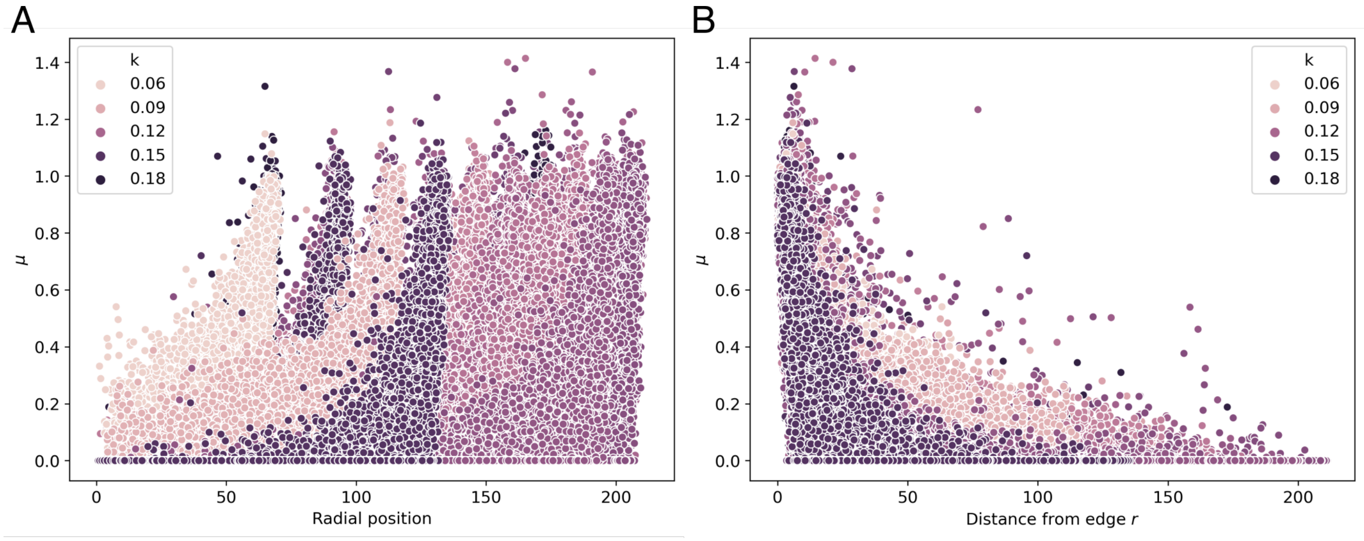


Figure 1: Growth rate profile at different colony sizes. **(A)** Distribution of individual bacteria growth rate at different $R_{max}$ or colony sizes ordered by radial position. **(B)** Distribution of individual bacteria growth rate at different $R_{max}$or colony sizes ordered by distance from edge $r$.

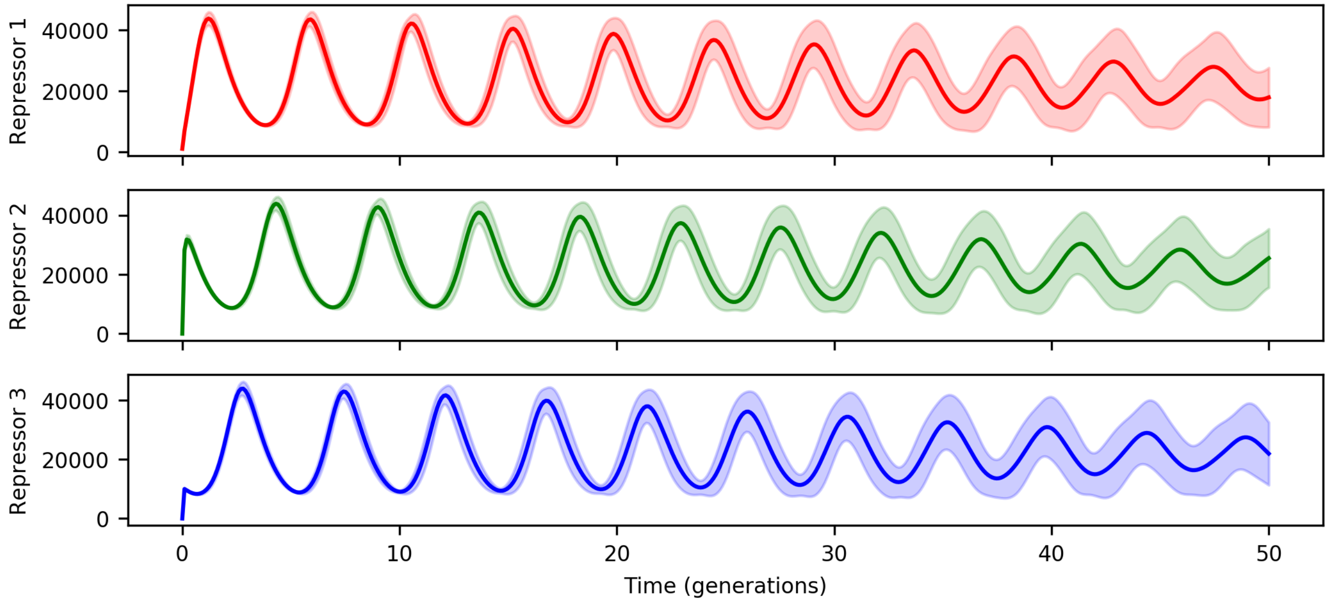


Figure 2: Stochastic simulation algorithm based on model of [1] using $K = 10^3$, $\lambda = 10^5$, $n = 2$, $b = 1$, and $N_0 = 100$. The mean and standard deviation of 100 simulations are shown.

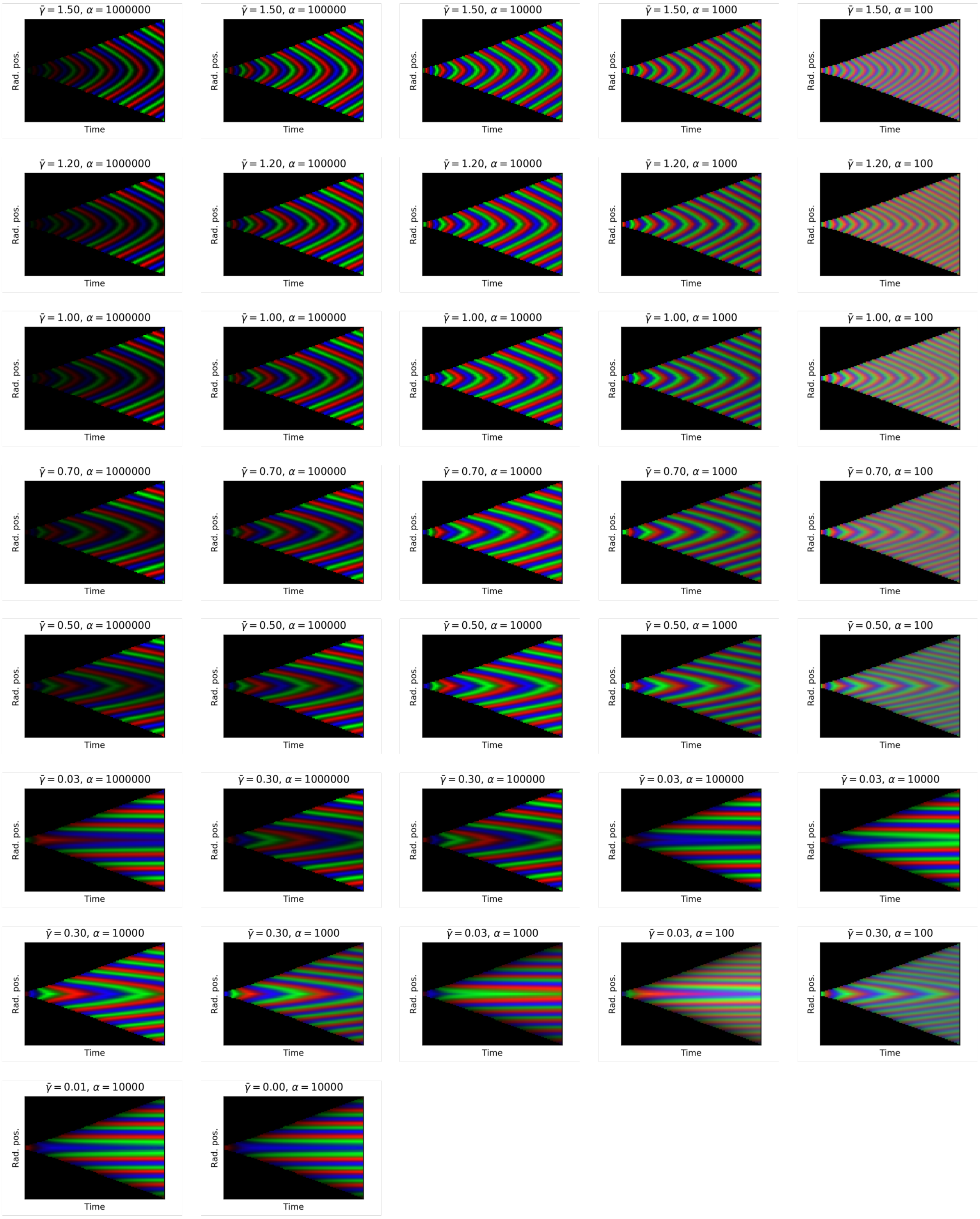


Figure 3: Kymographs computed from individual based model simulations of growing colonies from 1 to approximately 60,000 cells.

#### 6 Colonies

Here we show the four colonies analyzed in figure 5 of the main text, when composed of 60.000 cells. Figure 4 shows the colony with $\bar{\gamma}$=0.0 and $\alpha$=10,000. Figure 5 shows the colony with $\bar{\gamma}$=0.3 and $\alpha$=10,000. Figure 6 shows the colony with $\bar{\gamma}$=0.3 and $\alpha$=100. Figure 7 shows the colony with $\bar{\gamma}$=1.0 and $\alpha$=1000.

#### References

[1] Laurent Potvin-Trottier et al. "Synchronous long-term oscillations in a synthetic gene circuit". In: *Nature* 538.7626 (2016), pp. 514–517. DOI: 10.1038/nature19841.

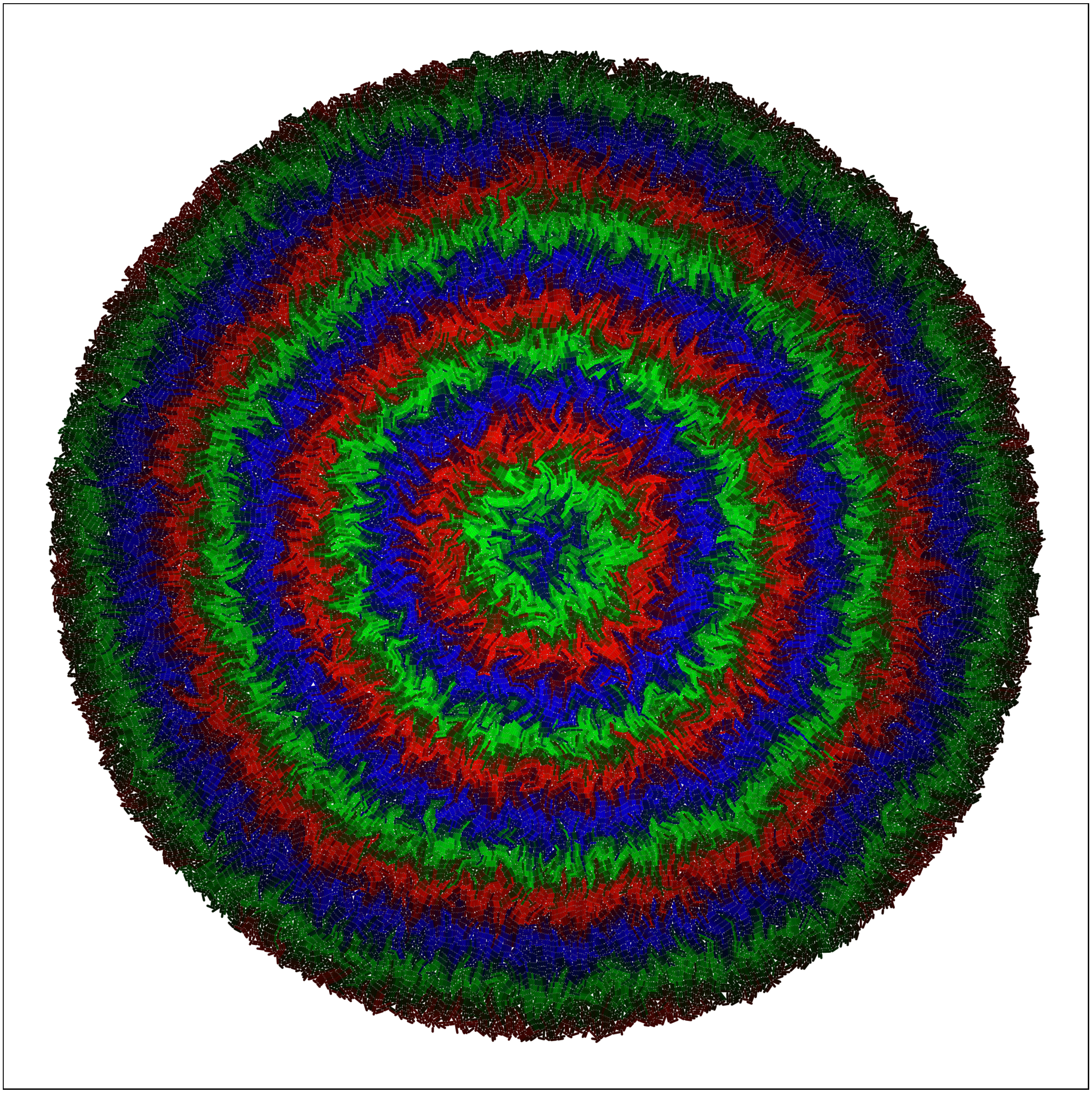

Figure 4: 60.000 cells. $\bar{\gamma}$=0 and $\alpha$=10,000.

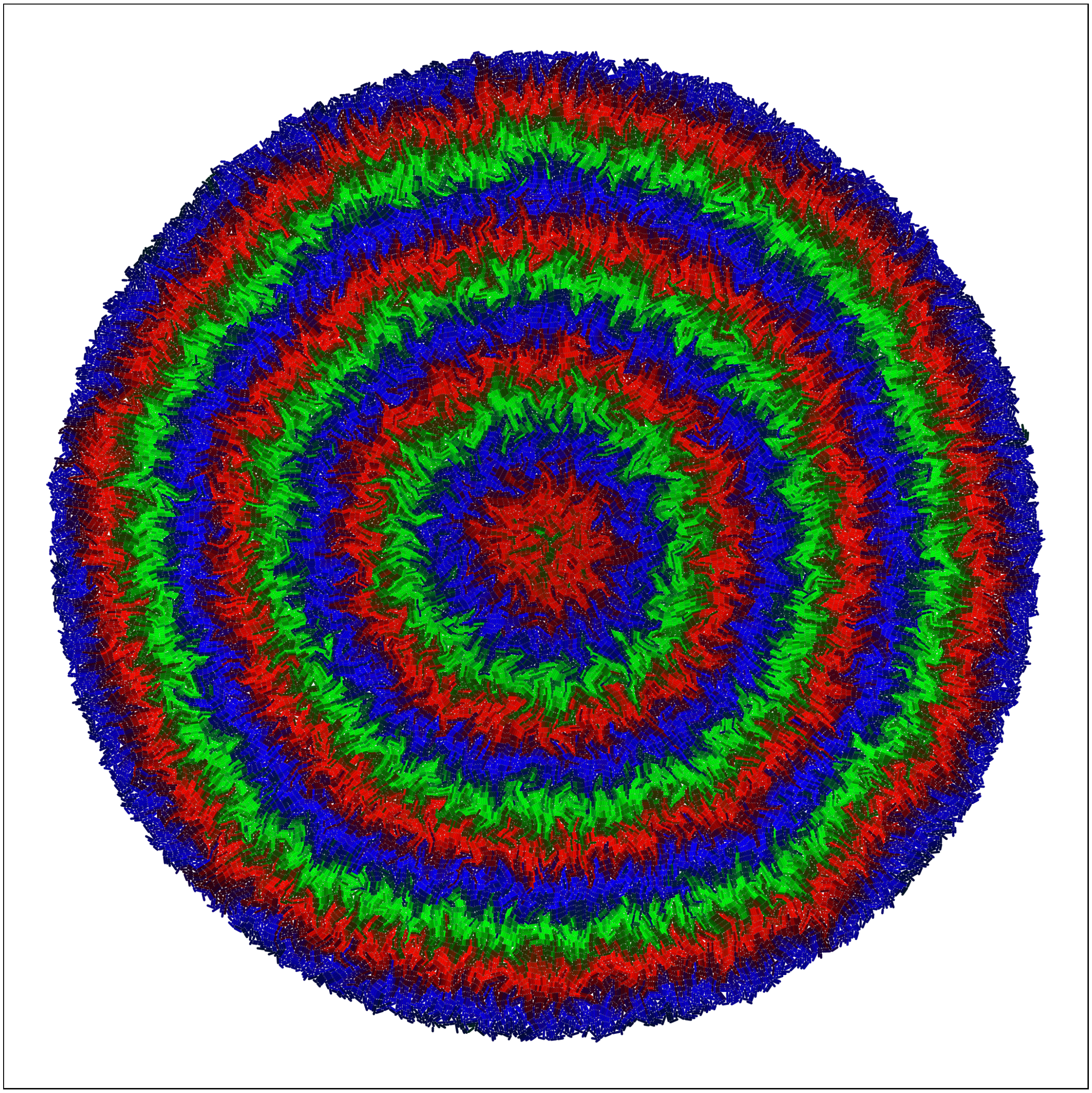

Figure 5: 60.000 cells. $\bar{\gamma}$=0.3 and $\alpha$=10,000.

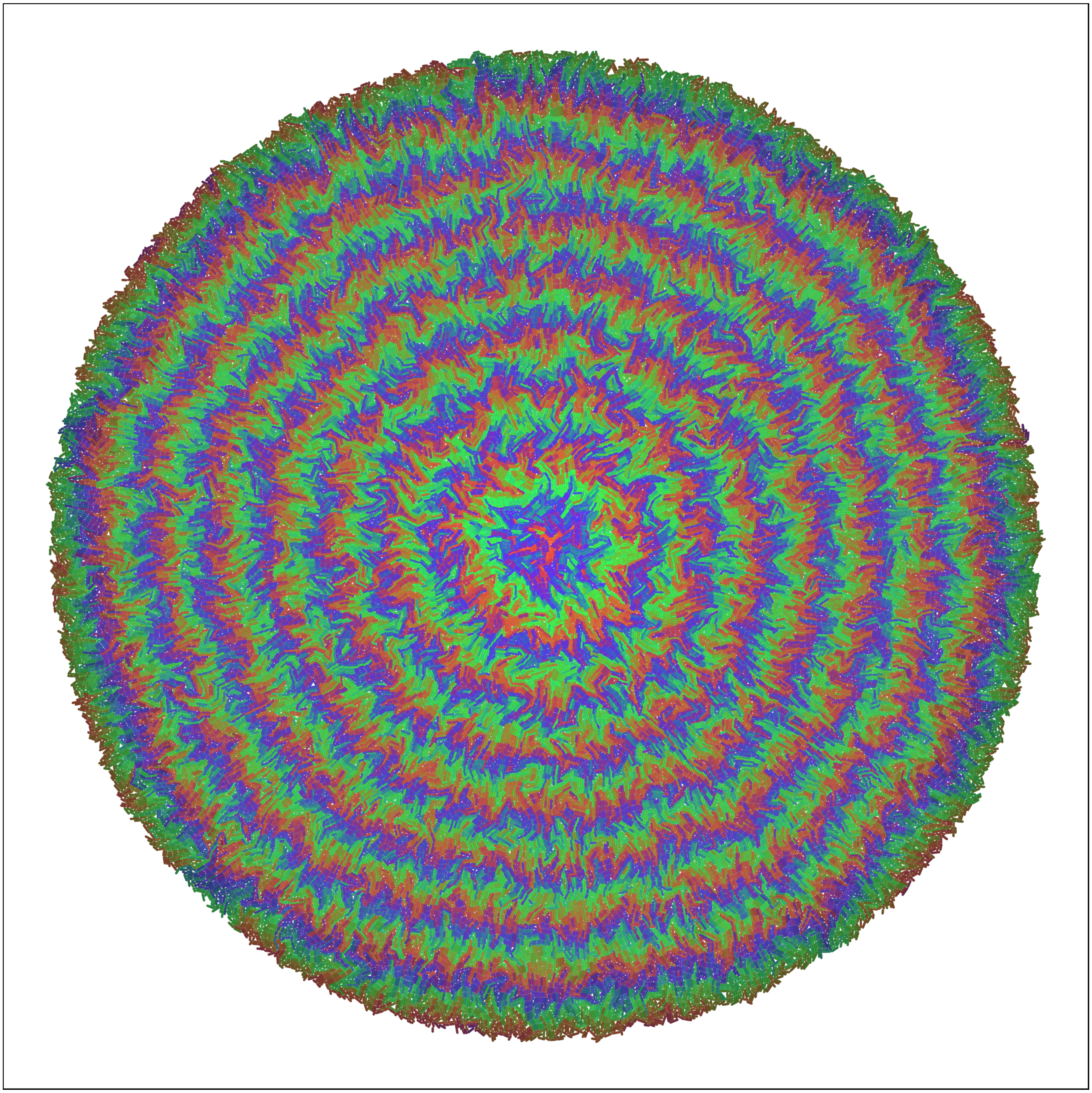

Figure 6: 60.000 cells. $\bar{\gamma}$=0.3 and $\alpha$=100.

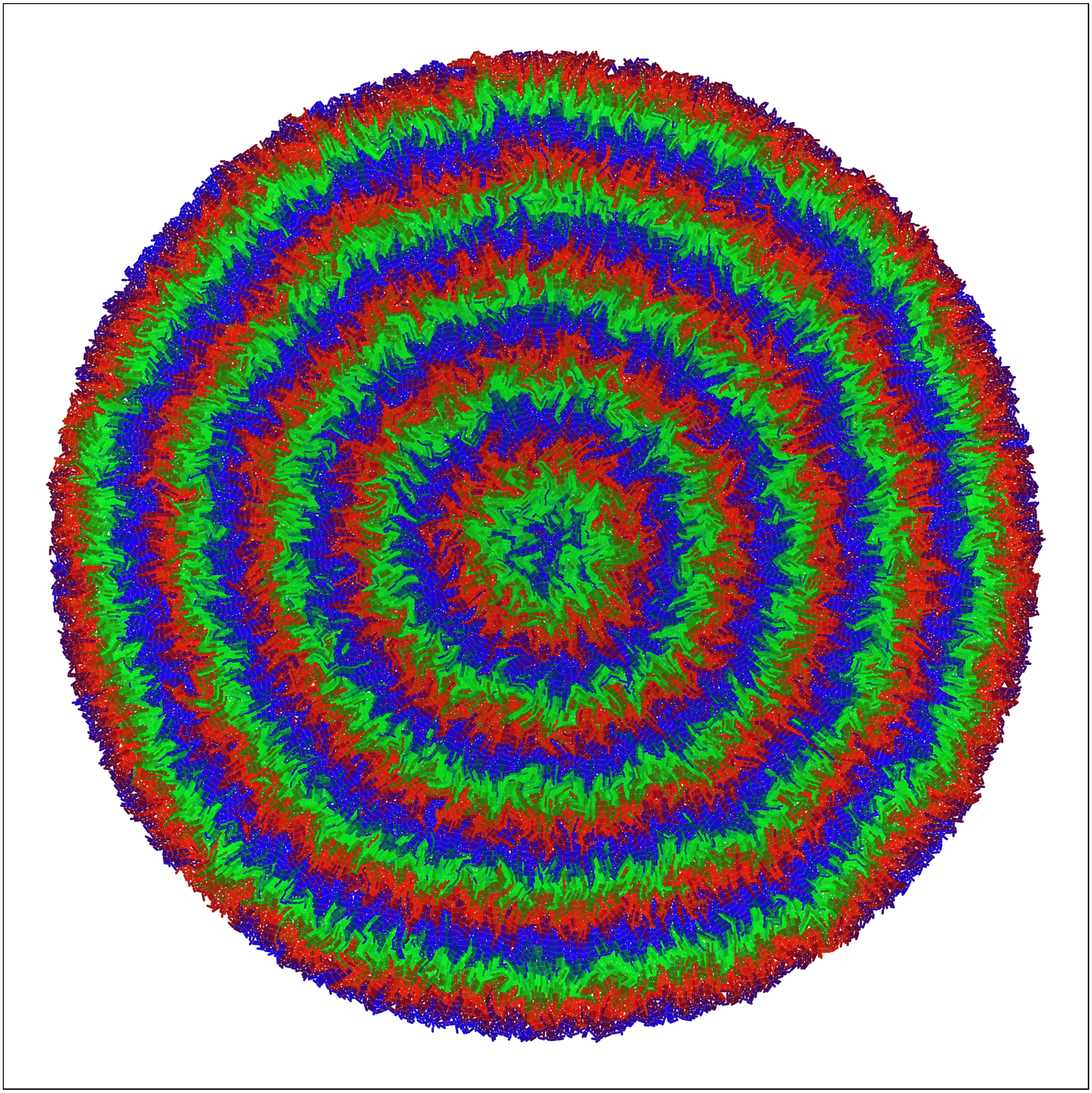

Figure 7: 60.000 cells. $\bar{\gamma}$=1 and $\alpha$=1000.
